# CRISPR screens in sister chromatid cohesion defective cells reveal PAXIP1-PAGR1 as regulator of chromatin association of cohesin

**DOI:** 10.1101/2022.12.23.521474

**Authors:** Janne J.M. van Schie, Klaas de Lint, Thom M. Molenaar, Macarena Moronta Gines, Jesper A. Balk, Martin A. Rooimans, Khashayar Roohollahi, Govind M. Pai, Lauri Borghuis, Anisha R. Ramadhin, Josephine C. Dorsman, Kerstin S. Wendt, Rob M.F. Wolthuis, Job de Lange

## Abstract

The cohesin complex regulates higher order chromosome architecture through maintaining sister chromatid cohesion and folding chromatin by active DNA loop extrusion. Impaired cohesin function underlies a heterogeneous group of genetic syndromes and is associated with cancer. Here, by using synthetic lethality CRISPR screens in isogenic human cell lines defective of specific cohesion regulators, we mapped the genetic dependencies induced by absence of DDX11 or ESCO2. The obtained high confidence synthetic lethality networks are strongly enriched for genes involved in DNA replication and mitosis and support the existence of parallel sister chromatid cohesion establishment pathways. Among the hits, we identified the chromatin binding, BRCT-domain containing protein PAXIP1 as a novel cohesin regulator. Depletion of PAXIP1 severely aggravated cohesion defects in ESCO2 defective cells, leading to mitotic cell death. PAXIP1 promoted the global chromatin association of cohesin, independent of DNA replication, a function that could not be explained by indirect effects of PAXIP1 on transcription or the DNA damage response. Cohesin regulation by PAXIP1 required its binding partner PAGR1 and a conserved FDF motif in PAGR1. Similar motifs were previously found in multiple cohesin regulators, including CTCF, to mediate physical interactions with cohesin. PAXIP1 co-localizes with cohesin on multiple genomic loci, including at active gene promoters and enhancers. Together, this study identifies the PAXIP1-PAGR1 complex as a novel regulator of cohesin occupancy on chromatin. Possibly, this role in cohesin regulation is also relevant for previously described functions of PAXIP1 in transcription, immune cell maturation and DNA repair.

## INTRODUCTION

The cohesin complex regulates structural genome organization, thereby contributing to critical cellular processes including transcription, DNA repair and chromosome segregation. The SMC1/SMC3 heterodimer, the kleisin subunit RAD21 and one SA subunit (SA1 or SA2) together form a circular structure that can physically tether DNA molecules ^1^. This occurs *in trans*, to facilitate chromosome segregation and homology directed DNA repair, and *in cis*, to create chromatin loops that can form topologically associating domains (TADs) and play roles in transcriptional regulation ^2^. Defective cohesin function and mutations in cohesin genes and regulators are associated with cancer ^3^ and underlie a group of rare developmental disorders, termed cohesinopathies ^4^.

The establishment of sister chromatid cohesion is tightly coupled to DNA replication ^5^. Cohesion establishment depends on the acetylation of SMC3 by ESCO2 during DNA replication to promote the association of Sororin, which counteracts cohesin dissociation by the cohesin release factor WAPL ^6-9^. In addition, several other replisome associated proteins are involved. Based on epistasis studies in yeast, these are suggested to function in two parallel pathways ^10, 11^. One pathway requires Chl1, Csm3-Tof1 and Ctf4 (DDX11, TIMELESS-TIPIN and AND-1 in human) and is believed to depend on cohesin complexes that were pre-loaded onto chromatin in G1 ^12^. The second pathway involves the alternative PCNA loader subunits Ctf18-Ctf8-Dcc1 (CHTF18-CHTF8-DSCC1 in human) and depends on *de novo* loading of cohesin during DNA replication by Scc2 (NIPBL in human) ^12^. To what extent these findings in yeast also apply to human cells, and which additional factors contribute to cohesion establishment, remains to be determined.

Although sister chromatid cohesion is essential for cellular proliferation, partial loss of cohesion can be tolerated to different degrees. Cell lines derived from patients with the cohesinopathies Warsaw Breakage Syndrome and Roberts Syndrome, caused by mutations in the DNA helicase DDX11 and the acetyltransferase ESCO2, respectively, are both characterized by premature loss of cohesion in metaphase. While these cells are viable, this phenotype creates specific vulnerabilities. Combined impairment of redundant cohesion establishment pathways is synthetically lethal due to enhanced cohesion defects beyond tolerable levels ^10, 11, 13-15^. Moreover, cohesion defects and inactivating mutations in cohesion related genes can sensitize cells to prolonged metaphase duration and drugs that induce DNA damage ^16-23^. Identification of the factors that determine cohesion proficiency may be clinically relevant, as they point at potential vulnerabilities of cohesion defective cancer cells ^22^.

Only a fraction of chromatin-bound cohesin maintains sister chromatid cohesion ^24, 25^, while cohesin also contributes to intra-chromosomal DNA-DNA contacts. Cohesin occupancy on chromatin is regulated by the dynamic loading, translocation and unloading of cohesin. The NIPBL-MAU2 heterodimer loads cohesin onto DNA, while WAPL promotes release of cohesin from chromatin. NIPBL-bound cohesin mediates active DNA loop extrusion to fold DNA fibers ^26, 27^, thereby contributing to 3D chromosome organization and *in cis* DNA-DNA contacts including enhancer-promotor loops involved in transcriptional regulation ^2^. CCCTC-binding factor (CTCF) can anchor cohesin translocation by interacting with cohesin via a conserved F/YxF motif, thereby enriching cohesin occupancy at CTCF sites and stabilizing cohesin onto chromatin by restricting WAPL binding ^28^. In addition, various other chromatin-binding proteins, including chromatin remodelers, are reported to interact with and enrich cohesin and/or its loader at specific chromatin sites ^29-36^. Thereby, spatiotemporal cohesin occupancy on chromatin is tightly coordinated with and influenced by chromatin dynamics.

In this study, we aimed to map the human genes involved in modulating sister chromatid cohesion and to find mechanisms of tolerance against partial sister chromatid cohesion loss. By using genome-wide CRISPR screens in DDX11 and ESCO2 defective human cell lines, we generated a network of synthetic lethal interactors of cohesion loss. Among our hits we identified the chromatin associated PAXIP1-PAGR1 complex as a novel regulator of cohesin association with chromatin.

## RESULTS

### Isogenic genome-wide CRISPR screens identify known and novel genetic interactors of DDX11 and ESCO2

To identify synthetic interactions in cells with impaired cohesion establishment, we used RPE1-hTert-TP53KO cells with inducible Cas9 ^37^ (hereafter referred to as RPE1-WT) to create isogenic cell lines with mutations in DDX11 and ESCO2. We generated a DDX11-KO cell line, but were unable to generate a viable ESCO2-KO cell line, similar as reported previously ^38^. Instead, we used a hypomorphic ESCO2 mutant with a mutation in the PDM-A domain (Fig. S1A). In line with previous studies ^39, 40^, this resulted in reduced ESCO2 protein levels that only become detectable after inhibition of the proteasome (Fig. S1B). The resulting DDX11-KO and ESCO2-mut cell lines show no detectable DDX11 and strongly reduced ESCO2 levels respectively (Fig. 1A) and display pronounced cohesion defects (Fig. 1B), confirming their functional impairment.

**Figure 1.**
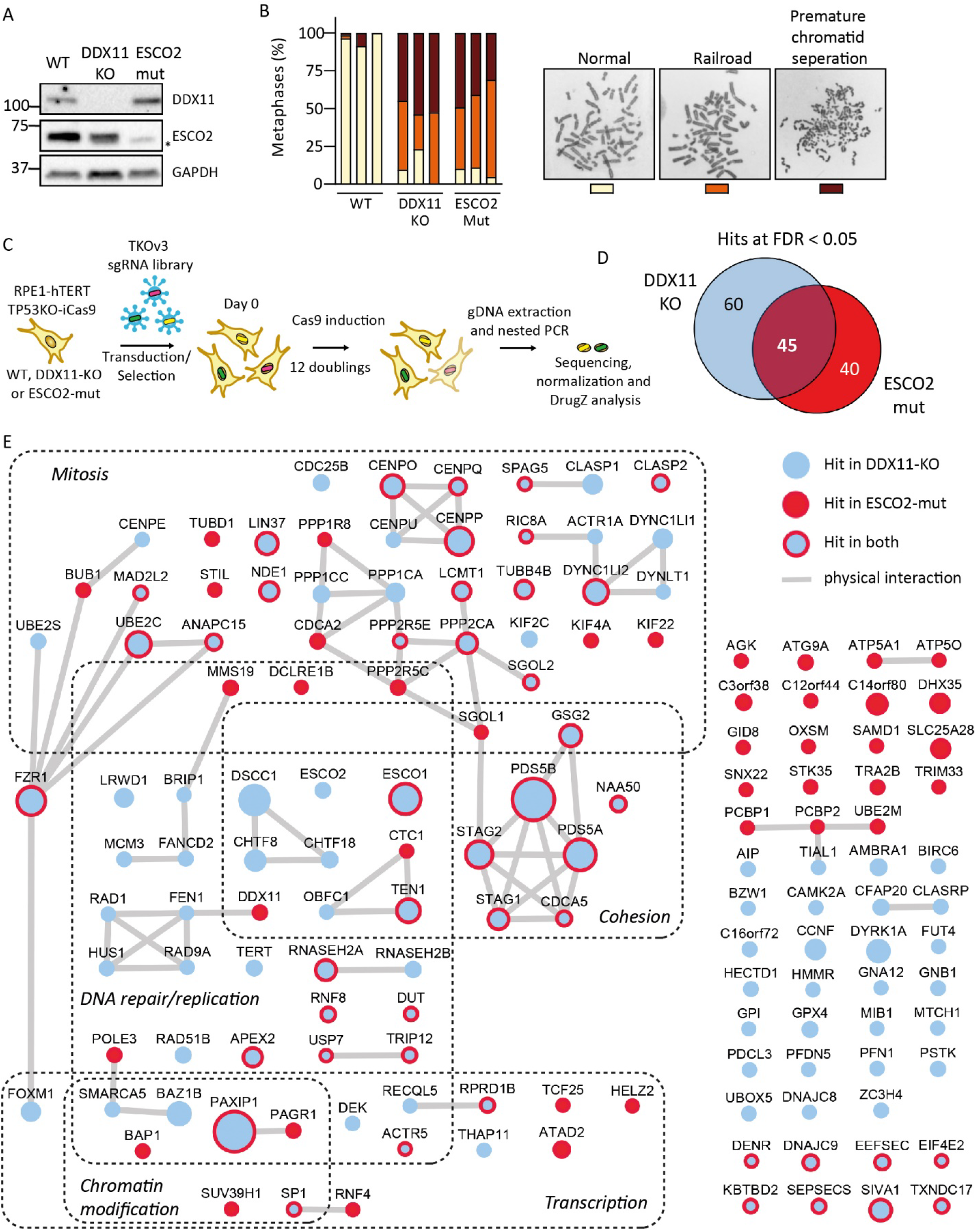
Genome-wide CRISPR screens identify genetic interactions with ESCO2 and DDX11. (**A**) Western blot of DDX11-KO and ESCO2-mutant cell lines. Asterisk indicates non-specific band. (**B**) Cohesion defect analysis of indicated cell lines. N=50 for each sample, three independent experiments are shown as separate bars. (**C**) Workflow of CRISPR screens, performed in triplicate. (**D**) Venn diagram of identified hits in both screens with FDR < 0.05 (DrugZ). (**E**) Network analysis of hits representing synthetically lethal genes with DDX11 (blue), ESCO2 (red) or both (blue-red). Edges indicate physical protein–protein interactions (String-db, evidence-based). Node size reflects significance, using the highest value from the two screens. Table S1 contains the raw data.

DDX11-KO, ESCO2-mut and WT cell lines were transduced with the genome-wide TKOv3 library ^41^ and cultured for 12 population doublings, followed by genomic DNA isolation and sequencing of sgRNA inserts (Fig. 1C). The results of the WT screen were published previously ^37^. After normalization based on t=0 counts, we computed WT versus DDX11-KO and WT versus ESCO2-mut gene-level depletion scores using DrugZ ^42^ (Table S1). With an FDR < 0.05 we identified 105 and 85 synthetic lethal interactions, respectively, of which 45 overlapped between the two screens (Fig. 1D). The hits are enriched in genes involved in sister chromatid cohesion, mitosis and DNA replication and repair (Fig. 1E and Fig. S1C). Both screens identified multiple previously described genetic dependencies, including synthetic lethality between DDX11 and ESCO2 ^13, 14^ and the ESCO2 paralog ESCO1, which is particularly essential in ESCO2-mut cells ^13, 15, 38^. Similar as reported in budding yeast and vertebrate cell lines ^10, 23, 43^, DDX11-KO cells showed increased dependency on CHTF18, CHTF8 and DSCC1, the specific subunits of the alternative replication factor C clamp loader CHTF18-RFC. Moreover, in line with the role of DDX11 in mitigating DNA replication stress ^17, 19, 44^, DDX11-KO cells were particularly sensitive to depletion of multiple DNA replication and repair factors, such as BRIP1 (FANCJ), FANCD2, RAD51B and the RAD9A-HUS1-RAD1 (9-1-1) complex. Together, our screens yield high confidence networks of genetic dependencies of ESCO2 and DDX11, providing a rich resource of the human cohesin regulatory network.

To validate the identified synthetic lethal interactions, we reconstituted the knockout cell lines with ectopic DDX11 or ESCO2 (Fig. S2A-B). Subsequently, we depleted a selection of hits using Cas9 induction and either synthetic crRNA transfections (Fig. 2) or viral sgRNA transductions (Fig. S2C) and assessed cell proliferation. This confirmed increased sensitivity to depletion of the cohesion factors ESCO1, ESCO2, DDX11, DSCC1, PDS5A and PDS5B. Remarkably, whereas ESCO2-mut cells are more sensitive to depletion of PDS5A than PDS5B, this is reversed in DDX11-KO cells, suggesting partially separate functions of the PDS5 homologs. In addition to known cohesion factors, the majority of other hits that we tested could also be validated, including PAXIP1, PAGR1, SIVA1, DYNC1Li2, BAZ1B, FZR1, RNF8, CENPP and CENPO, further confirming the high confidence of hits found in our screens.

**Figure 2.**
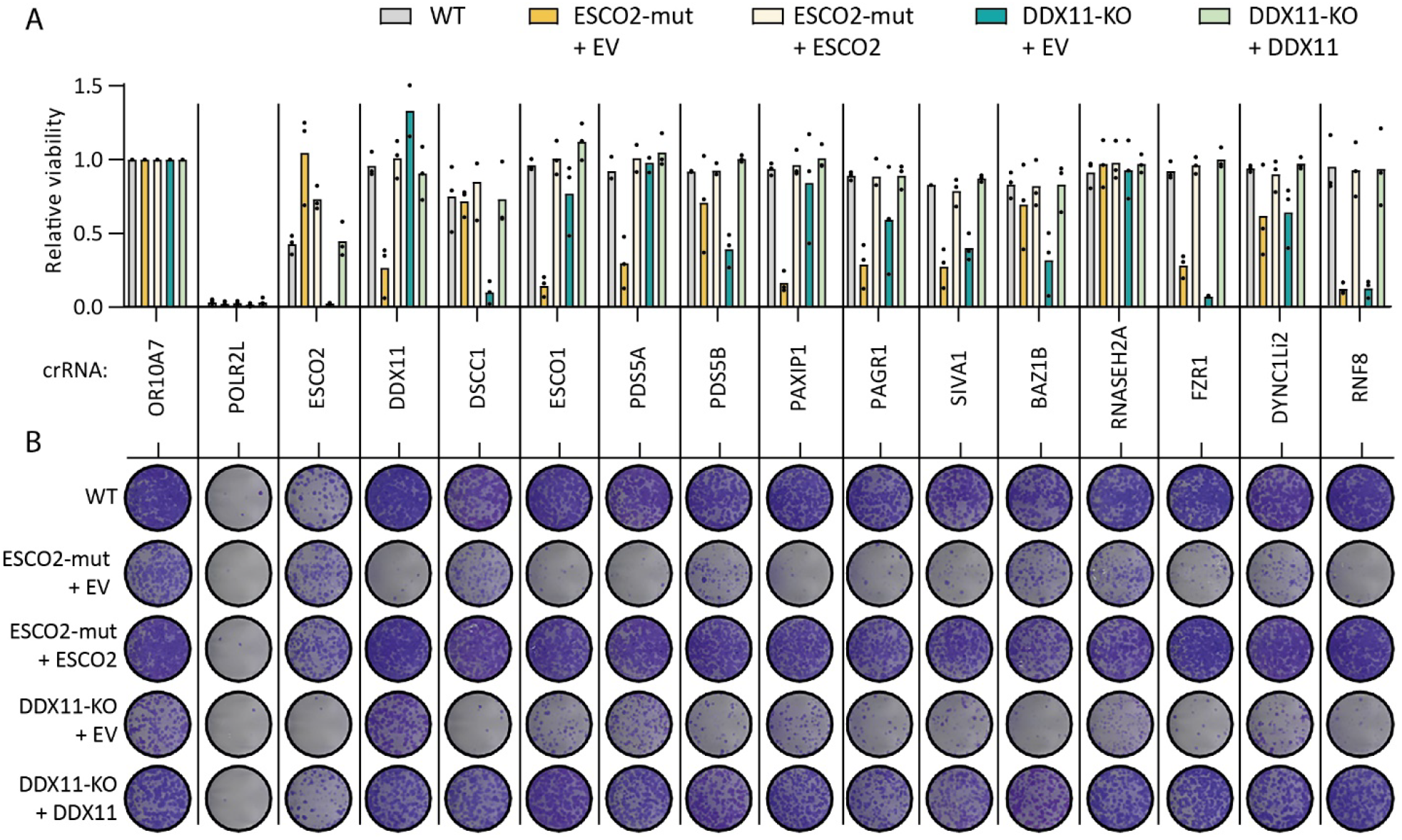
Hit validation in ESCO2-mut and DDX11-KO cells. (**A**) Viability of WT, ESCO2-mut and DDX11-KO cells assessed by a CellTiter-Blue assay 7 days after crRNA:tracrRNA transfection and accompanying Cas9 induction. Viability is normalized to cells transfected with crOR10A7, a nonessential gene. POLR2L is used as a common essential control. Dots represent the mean of three technical replicates; bars indicate the mean of three independent experiments. (**B**) Clonogenic survival assay 10 days after crRNA:tracrRNA transfection in indicated cells and accompanying Cas9 induction.

As a representative mitosis-associated hit, we further analyzed the non-catalytic microtubule motor protein subunit DYNC1Li2. Knockdown of DDX11 and ESCO2 caused elevated mitotic fractions and cohesion loss in DYNC1Li2-KO cells compared to WT cells (Fig. S3). This indicates that DYNC1Li2 is redundant for sister chromatid cohesion but becomes critical to mitigate further loss of cohesion in cohesion defective cells, possibly due to its role in metaphase duration, similar to what was previously shown for impaired APC/C function ^22^.

### PAXIP1 promotes the chromatin association of cohesin

Among the top validated hits in the ESCO2 screen was PAXIP1, a chromatin binding protein with described functions in transcriptional regulation, the DNA damage response and immune cell maturation ^45-47^, but no described role in cohesin biology. In order to study PAXIP1 in more detail, we made clonal PAXIP1-KO RPE1 cells (Fig. S4A). Considering the role of PAXIP1 as a transcriptional regulator, we performed RNA sequencing (Fig. S4B-C and Table S2). While the expression of 664 genes was significantly changed in PAXIP1-KO cells (357 genes downregulated and 307 upregulated in PAXIP1-KO), no changes were observed in cohesin related genes or other hits from the CRISPR screen, suggesting that an altered transcriptional profile does not directly explain the observed synthetic lethality of PAXIP1 and ESCO2. An alternative explanation could lie in the role of PAXIP1 in the DNA damage response ^48-50^. However, we observed no increased γH2AX signaling in interphase (Fig. S4D). Of note, DNA damage could be observed upon PAXIP1 depletion in a fraction of ESCO2-mut cells with disrupted nuclear integrity (Fig. S4D), which is likely a secondary effect linked to mitotic catastrophe ^51^.

Interrogation of the DepMap database (https://depmap.org) revealed a remarkable correlation of essentiality of PAXIP1 with the cohesin subunits SMC3 and STAG2 and the loaders NIPBL and MAU2 (Fig. S4E). Since co-essentiality implies shared biological function ^52^, this led us to hypothesize that PAXIP1 may directly influence cohesin function. Although sister chromatid cohesion and cell cycle distribution were unaffected in PAXIP1-KO clones (Fig. S4F-G), we observed an aggravation of cohesion loss (Fig. 3A) and accumulation of mitotic cells (Fig. 3B) upon PAXIP1 depletion in ESCO2-mut cells, suggesting mitotic cell death resulting from detrimental cohesion defects. Strikingly, we detected reduced levels of chromatin-associated cohesin by RAD21 immunofluorescence in PAXIP1-KO cells (Fig. 3C) and upon PAXIP1 knockdown (Fig. S4H). This was confirmed by immunoblotting RAD21, SMC3 and NIPBL in chromatin-bound protein fractions (Fig. 3D). Chromatin-bound cohesin was reduced in both G1 and G2 synchronized cells (Fig. 3D and Fig. S4I-J), suggesting that this function of PAXIP1 is independent of DNA replication. Importantly, in line with the RNA-seq data, soluble and total levels of cohesin proteins were similar in WT and PAXIP1-KO cells (Fig. 3D and Fig. S4J). Together, these results suggest that PAXIP1 directly influences the chromatin association of cohesin throughout the cell cycle.

**Figure 3.**
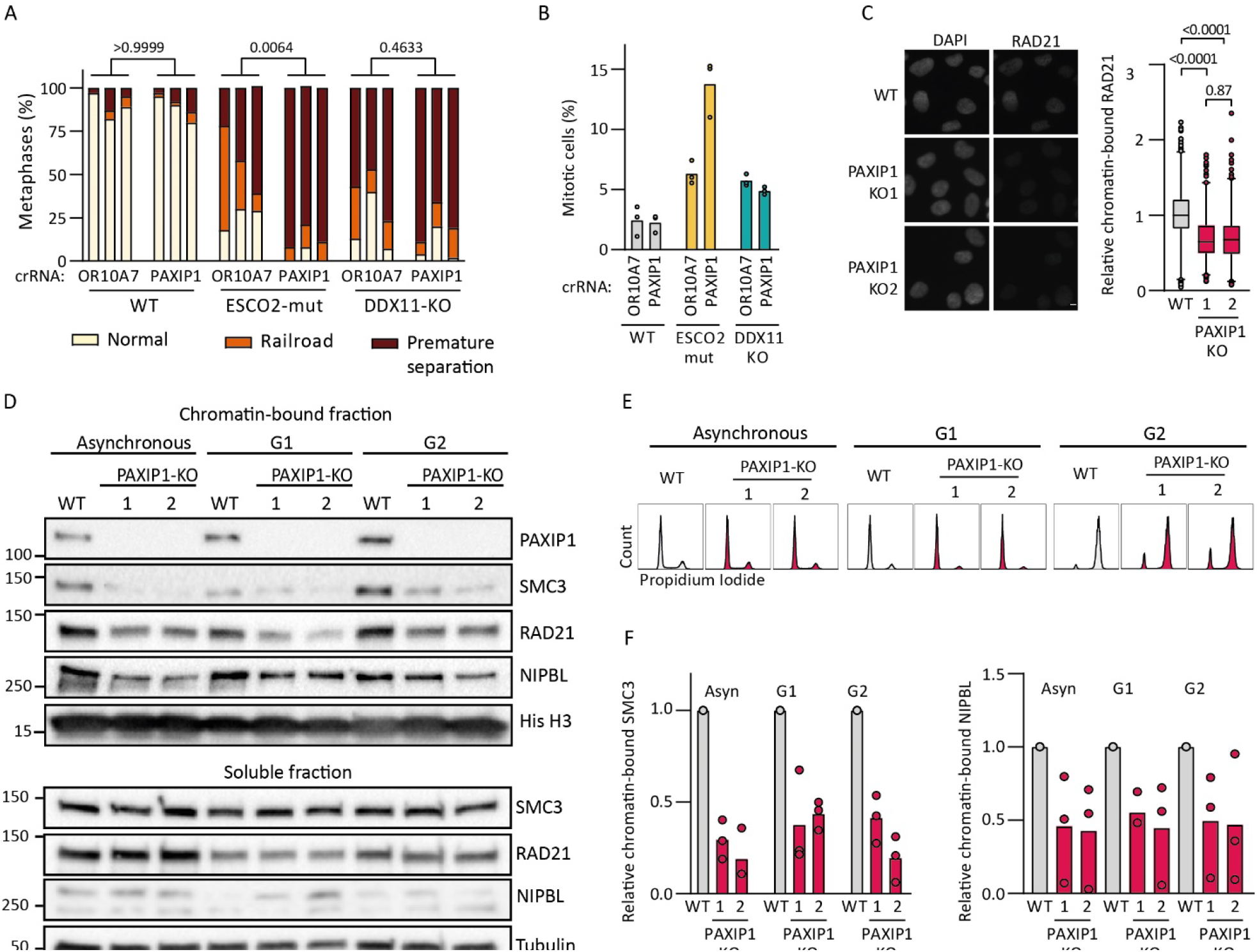
PAXIP1 promotes chromatin association of cohesin throughout the cell cycle. (**A**) Cohesion defect analysis two days after crPAXIP1 transfection. N=50 for each sample, three independent experiments are shown as separate bars. P-values were calculated by a one-way ANOVA comparing the frequency of premature chromatid separation per condition. (**B**) Percentage of mitotic cells two days after crPAXIP1 transfection assessed by flow cytometry of p-Histone H3-S10 stained cells from three independent experiments. (**C**) Chromatin-bound cohesin levels of WT, PAXIP1-KO1 and PAXIP1-KO2 cells assessed by RAD21 immunofluorescence of pre-extracted cells. RAD21 intensity was quantified for at least 205 cells per sample in three independent experiments. P-values were calculated by a one-way ANOVA. Box represents 25%-75% and median, whiskers represent 1-99 percentile. Scale bar represents 5 µM. (**D**) Western blot of chromatin-bound and soluble protein fractions in WT and PAXIP1-KO cells in asynchronous cells (Asyn) or synchronized cells, in G1 by 10µM Palbociclib or in G2 by 10µM RO-3306. (**E**) Flow cytometry control of the synchronization in D. (**F**) Quantification of chromatin-bound SMC3 and NIPBL relative to WT from three independent experiments.

### PAXIP1 function in cohesin regulation depends on the interaction with PAGR1

PAXIP1 can physically associate with the histone methyltransferases KMT2C and KMT2D, the DNA damage response protein TP53BP1, and PAXIP1 associated glutamate-rich protein (PAGR1) ^46, 47, 53-55^. The ESCO2-mut CRISPR screen identified PAGR1, but not TP53BP1, KMT2C/D or any of the accessory subunits of the KMT2C/D methyltransferase complex (WDR5, RBBP5, ASH2L, DPY30, KDM6A) (Fig. 1E and Table S1). Co-depletion of KMT2C and KMT2D, which may in part be functionally redundant ^56^, also did not impair growth of ESCO2-mut cells (Fig. S5A-B), and chromatin-bound cohesin levels were not affected in KMT2C/D double KO (dKO) clones (Fig. S5C-F). Moreover, HCT116-KMT2D-KO cells ^57^, which already lack KMT2C ^58^, did not show a clear decrease in chromatin-bound cohesin (Fig. S5G-H), further suggesting that KMT2C/D is not involved in regulating cohesin levels on chromatin.

Synthetic lethality of PAGR1 was validated by crRNA transfections, which resulted in increased lethality, cohesion defects and an elevated mitotic fraction in ESCO2-mut cells (Fig. S6A-C), similar as observed upon PAXIP1 depletion (Fig. 3A-B). Next, we generated two PAGR1-KO clones using a crRNA targeting PAGR1 in its C terminal region (Fig. S6D). Although truncated protein products were still present (Fig. 4A), PAGR1 protein was functionally impaired, illustrated by destabilized PAXIP1 protein levels as previously reported ^46^. Similar to PAXIP1-KO cells, PAGR1-KO cells showed decreased chromatin bound cohesin levels (Fig. 4B-C) without causing cohesion defects in metaphase (Fig. S6E).

**Figure 4.**
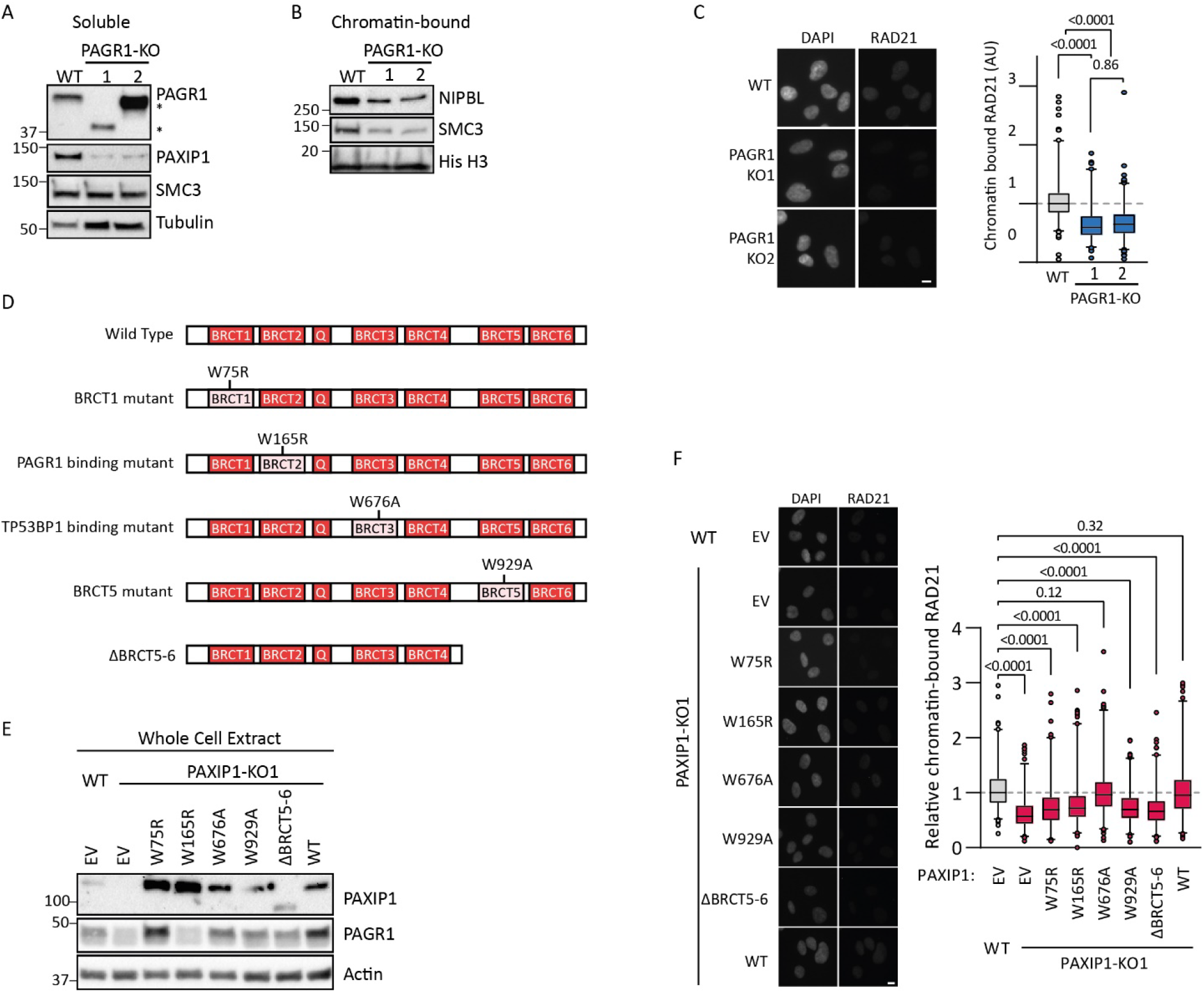
The role of PAXIP1 in maintaining chromatin-bound cohesin depends on the interaction with PAGR1. (**A**) Western blot of soluble protein fractions of WT and PAGR1-KOs cells. Asterisks indicate truncated proteins in PAGR1-KO clones, presumably resulting from a premature stop codon. (**B**) Western blot of chromatin bound protein fractions of WT and PAGR1-KOs cells. (**C**) RAD21 immunofluorescence in pre-extracted PAGR1-KO cells. Intensity was quantified from three independent experiments with at least 155 cells per condition. P-values were calculated by a one-way ANOVA. Box represents 25%-75% and median, whiskers represent 1-99 percentile. Representative images are shown on the left. Scale bar represents 10 µm. (**D**) Schematic representation of PAXIP1 expression constructs (BRCT = BRCA1 C-Terminus domain; Q = Glutamine-rich domain). (**E**) Western blot of whole cell extract of WT and PAXIP1-KO cells, stably transduced with PAXIP1 constructs described in 4D. (**F**) RAD21 immunofluorescence in pre-extracted in PAXIP1 mutant cells. Intensity was quantified from three independent experiments with at least 105 cells per sample. P-values were calculated by a one-way ANOVA. Box represents 25%-75% and median, whiskers represent 1-99 percentile. Representative images are shown on the left. Scale bar represents 10 µm.

PAXIP1 has six BRCT (BRCA1 C-Terminus) domains that mediate the association with different interaction partners ^46^. To investigate which BRCT domains of PAXIP1 contribute to the chromatin association of cohesin, we reconstituted PAXIP1-KO cells with WT or mutant forms of PAXIP1 (Fig. 4D). W75R is a BRCT1 mutant that does not abrogate PAGR1 binding but impairs the PAXIP1-PAGR1 sub-complex function, W165R perturbs the interaction with PAGR1, W676A with TP53BP1, and W929A and ΔBRCT5-6 were both reported to perturb the interaction of PAXIP1 with KMT2C/D ^46, 55^. PAXIP1 mutants were expressed at levels similar to or higher than endogenous PAXIP1 (Fig. 4E). This restored PAGR1 protein levels in PAXIP1-KO cells, except for the W165R mutant, in line with the reciprocal stabilization of PAXIP1 and PAGR1 following interaction ^46^. While PAXIP1-WT and W676A efficiently restored the level of chromatin-bound cohesin, no rescue was observed upon overexpression of W75R, W165R, W929A and ΔBRCT5-6 (Fig. 4F). This suggests that the role of PAXIP1 in promoting cohesin’s chromatin association depends on BRCT1, BRCT2, BRCT5 and BRCT6, but not on BRCT3. The fact that the W929A and ΔBRCT5-6 could not rescue the effects of PAXIP1-KO may suggest that BRCT5-6 has roles other than binding to KMT2C/D.

### Genetic interaction of PAXIP1 with the NIPBL-MAU2 cohesin loader complex

To characterize the genetic interactions of PAXIP1, we performed a genome-wide CRISPR screen in PAXIP1-KO cells. Among the 13 synthetically lethal hits at FDR < 0.1 were the cohesin factors STAG2, PDS5B and MAU2, further implicating PAXIP1 in cohesin regulation (Fig. 5A). Note that although ESCO2 depletion moderately further reduced viability of PAXIP1-KO cells (Fig. S7A), ESCO2 was not detected as a hit in the CRISPR screen. This could relate to the critical role of ESCO2 in cellular proliferation in both PAXIP1-KO as well as WT cells or to crRNA specific effects. Intrigued by the identification of a synthetic lethal interaction between PAXIP1 and MAU2, we decided to further investigate the relationship between the cohesin loader complex NIPBL-MAU2 and PAXIP1. Transfection of crMAU2 or crNIPBL resulted in proliferation defects, particularly in PAXIP1-KO cells (Fig. S7A-B). PAXIP1 loss further aggravated the decrease in chromatin-bound cohesin upon depletion of MAU2 or NIPBL (Fig. 5B-C). We then constructed stable MAU2-KO cells and subsequently made three MAU2-PAXIP1-dKO clones (Fig. S7C). MAU2 loss resulted in destabilization of NIPBL, in line with previous reports ^59, 60^ (Fig. 5D) and in slower proliferation compared to WT cells, which was further reduced in MAU2-PAXIP1-dKOs (Fig. 5E). Furthermore, MAU2-KO cells had lower levels of chromatin-bound cohesin compared to WT and PAXIP1-KO cells, which were slightly further reduced in MAU2-PAXIP1-dKOs (Fig. 5F). Possibly, PAXIP1 loss further diminishes the residual NIPBL-dependent cohesin loading activity in MAU2-KOs ^59, 60^. Alternatively, rather than facilitating cohesin loading, these observations may point at a role for PAXIP1 in stabilizing cohesin on chromatin.

**Figure 5.**
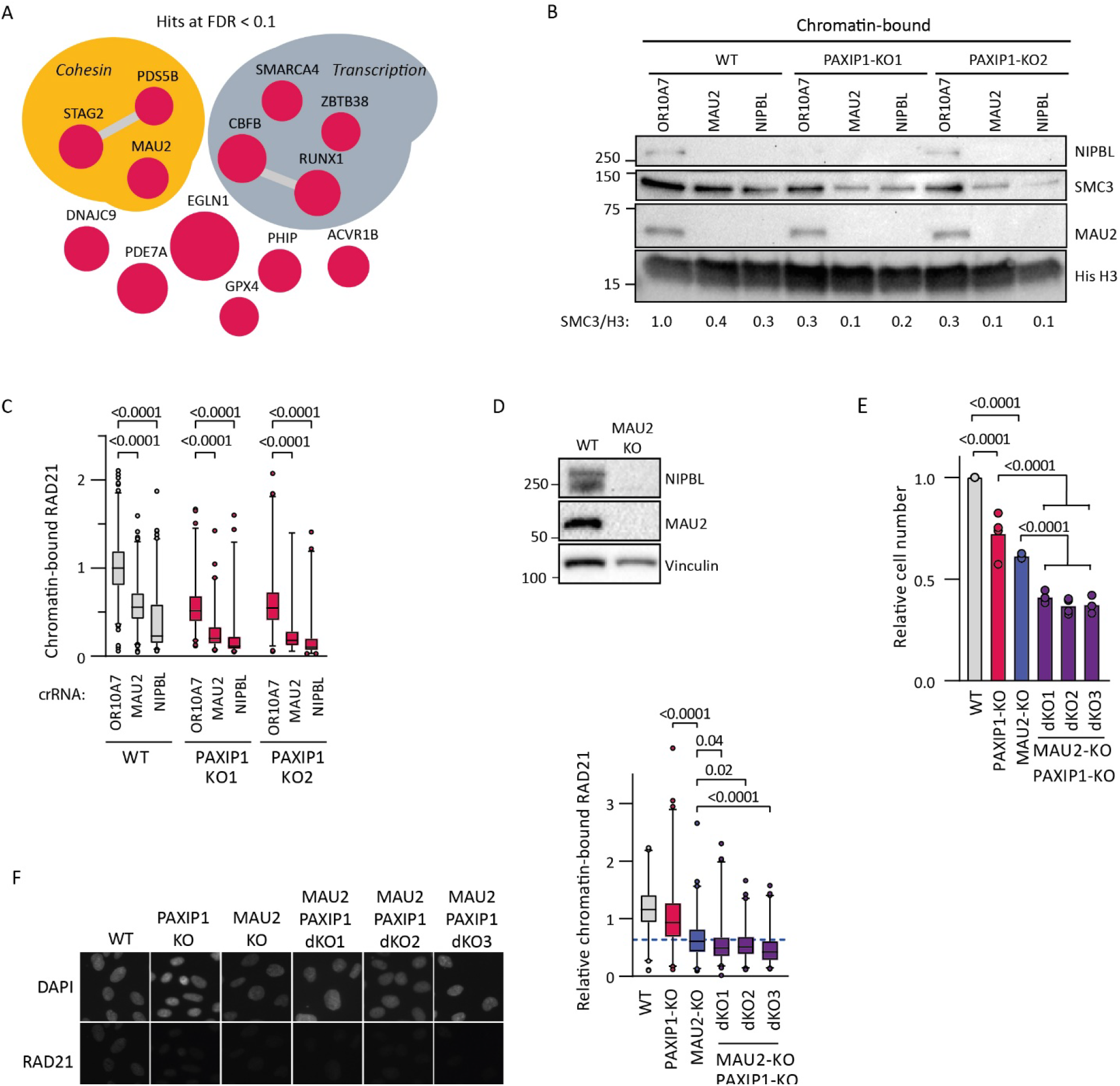
The genetic interaction between PAXIP1 and the cohesin loader NIPBL-MAU2. (**A**) Network of hits from a genome-wide CRISPR screen in PAXIP1-KO cells using FDR < 0.1. Edges indicate physical protein–protein interactions (String-db, evidence-based), node size reflects significance. Raw data in Table S1. (**B**) Western blot of chromatin-bound protein fractions of WT and PAXIP1-KOs after transfection with indicated crRNAs. SMC3 band intensities were quantified using Image Lab relative to WT crOR10A7 levels, corrected for histone H3 and are depicted below the blot. (**C**) WT and PAXIP1-KO cells were transfected with indicated crRNA’s and chromatin-bound cohesin was assessed by RAD21 immunofluorescence after pre-extraction. At least 88 cells were scored per condition in two independent experiments. P-values were calculated by a one-way ANOVA. Box represents 25%-75% and median, whiskers represent 1-99 percentile. (**D**) Western blot of whole cell extract from WT and MAU2-KO cells. (**E**) Cell counts relative to WT of indicated cells lines after four days proliferation. Dots represent the mean of two technical replicates; bars indicate the mean of four independent experiments. (**F**) Chromatin-bound cohesin assessed by RAD21 immunofluorescence after pre-extraction of PAXIP1-KO, MAU2-KO and PAXIP1-MAU2-doubleKO cell lines. At least 100 cells were scored per condition in three independent experiments. P-values were calculated by a one-way ANOVA. Box represents 25%-75% and median, whiskers represent 1-99 percentile. Scale bar represents 10 µm.

### A conserved FDF motif in PAGR1 promotes cohesin occupancy on chromatin

Interestingly, we discovered that PAGR1 contains a FDFDD motif, which is conserved in vertebrates (Fig. 6A). Similar F/YxF motifs (consensus [PFCAVIYL][FY][GDEN]F.(0,1)[DANE].(0,1)[DE]) have previously been found in several proteins, including CTCF, WAPL and MCM3, and were reported to mediate interactions at the STAG1/2-RAD21 interface and regulate cohesin dynamics on chromatin ^28, 61^. To test the relevance of this motif in PAGR1, we complemented PAGR1-KO1 with WT or mutant (ADA) FLAG-tagged PAGR1 (Fig. 6B). Both WT and mutant PAGR1 could stabilize PAXIP1 and maintained interaction with PAXIP1 and chromatin (Fig. 6B-D). However, unlike PAGR1-WT, the ADA mutant was unable to restore chromatin-bound cohesin levels (Fig. 6C, E-F). In conclusion, cohesin occupancy on chromatin depends on the FDF motif in PAGR1.

**Figure 6.**
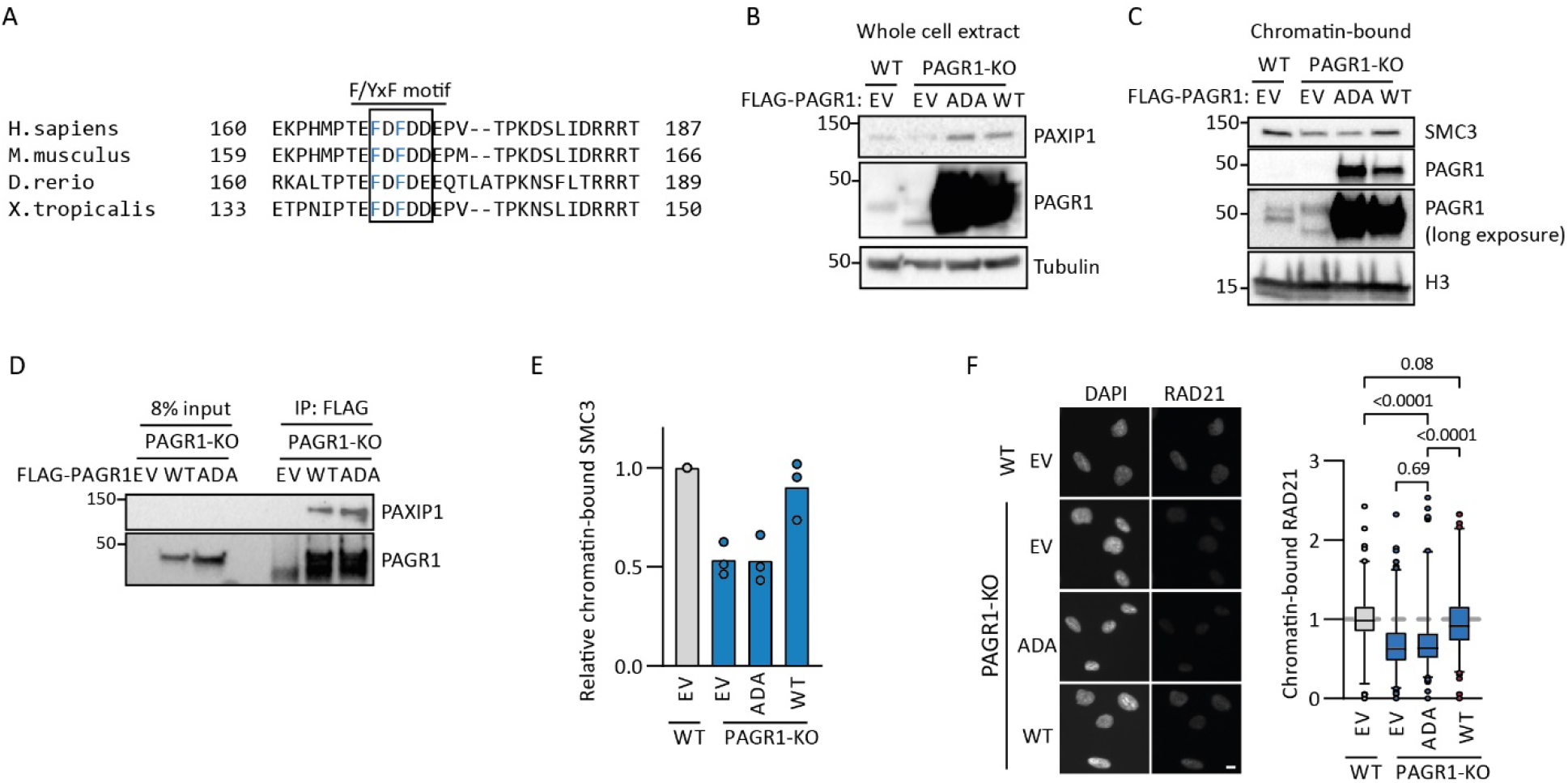
A conserved FDF motif in PAGR1 promotes cohesin occupancy on chromatin. (**A**) Sequence alignment of PAGR1 at the FDF motif. Box indicates the consensus FDFDD sequence, blue amino acids indicate phenylalanines mutated to alanines in the ADA mutant. (**B**,**C**) PAGR1-KO cells were stably transduced with empty vector (EV), FLAG-tagged WT or mutant (ADA) PAGR1. Whole cell extract (**B**) and chromatin-bound protein fractions (**C**) were assessed by western blot. (**D**) Flag-PAGR1 was immunoprecipitated from indicated cell lines followed by western blot analysis. (**E**) SMC3 band intensities were quantified from three independent western blots using Image Lab relative to WT-EV levels, corrected for histone H3. (**F**) RAD21 intensity was quantified in indicated cell lines from three independent experiments with at least 142 cells per sample. P-values were calculated by a one-way ANOVA. Box represents 25%-75% and median, whiskers represent 1-99 percentile. Scale bar represents 10 µm.

### PAXIP1 interacts and co-localizes with cohesin on chromatin at active promoters and enhancers

Since F/YxF motifs have been linked to cohesin binding ^28^, this may suggest PAXIP1-PAGR1 physically interacts with cohesin. In line, mass spectrometry of PAXIP1 co-precipitating proteins previously revealed SMC1 in HEK293T cells ^54^ and NIPBL in HeLa cells ^62^. Using co-immunoprecipitation experiments, we could indeed detect a physical interaction of endogenous PAXIP1 with SMC3, SA1, NIPBL and MAU2 in HEK293T cells (Fig. 7A) as well as of Venus-tagged PAXIP1 with RAD21 in RPE1 cells (Fig. 7B-C).

**Figure 7.**
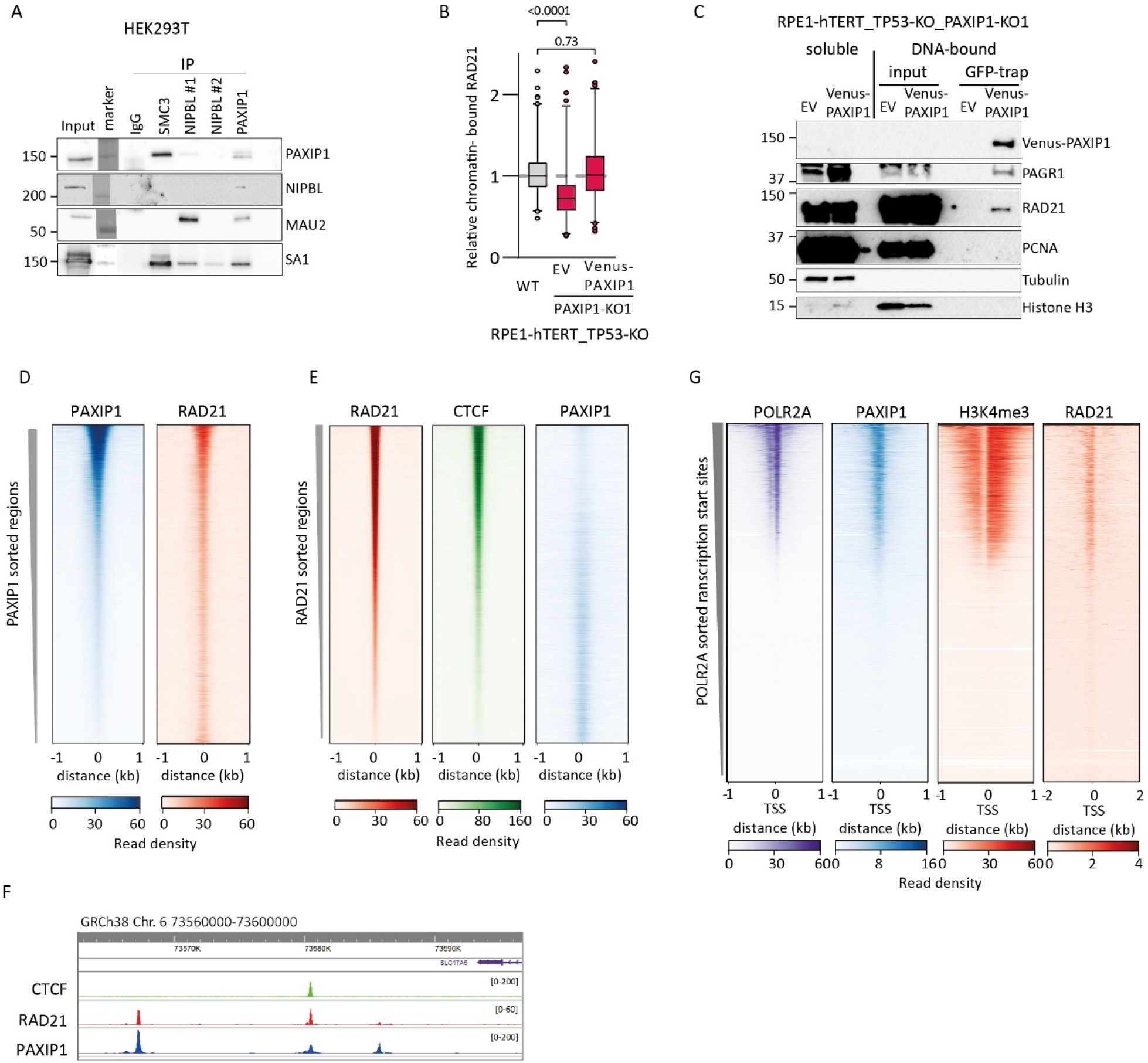
PAXIP1 interacts and co-localizes with cohesin on chromatin. (**A**) Co-immunoprecipitation using SMC3, NIPBL, PAXIP1 or IgG (control) antibodies in HEK293T cells, followed by western blot. (**B**) Chromatin-bound cohesin levels relative to WT of PAXIP1-KO1 RPE1 cells with empty vector (EV) or Venus-PAXIP1. At least 217 cells were scored per condition in two independent experiments. P-values were calculated by a one-way ANOVA. Box represents 25%-75% and median, whiskers represent 1-99 percentile. (**C**) Co-immunoprecipitation of Venus-PAXIP1 using GFP-trap beads in chromatin-bound protein fractions of RPE1 cells, followed by western blot. (**D**) Heatmaps of ENCODE ChIP-seq data for HepG2 cells sorted on PAXIP1 enriched regions. Regions are centered on PAXIP1 peaks -/+1kb. (**E**) Heatmaps of ENCODE ChIP-seq data for HepG2 cells sorted on RAD21 enriched regions. Regions are centered on RAD21 peaks -/+1kb. (**F**) Genome browser screenshot of ChIP-seq signals for RAD21, CTCF, and PAXIP1 in HepG2 cells. (**G**) Heatmaps of ENCODE ChIP-seq data for HepG2 cells sorted on transcription start sites (TSSs) enriched for POLR2A. Regions are centered on TSSs -/+1kb.

To determine if PAXIP1 localizes to cohesin-bound genomic regions, we mined publicly available ChIP-seq data for the human hepatocellular carcinoma cell line HepG2 from ENCODE ^63, 64^. This revealed that PAXIP1 occupied genomic sites that are enriched in RAD21 binding (Fig. 7D). As expected, sites with the strongest RAD21 signal co-localized with CTCF (Fig. 7E). Interestingly, PAXIP1 particularly occupied sites with a weaker CTCF signal (Fig. 7E-F), suggesting that PAXIP1 and cohesin preferentially co-localize at genomic loci that are less frequently bound by CTCF. To identify the chromosomal sites at which PAXIP1 and RAD21 co-localize, we examined promoters and enhancers. Previous reports described localization of PAXIP1 to promoters ^45, 65^. In line with this, we found that PAXIP1 is enriched at transcription start sites (TSSs) enriched for RNA Polymerase II (POLR2A) and H3K4me3, indicative of active promoters and co-localizes with RAD21 at these sites (Fig. 7G). In addition, PAXIP1 and RAD21 co-localize at active enhancers, defined by enrichment of the histone acetyltransferase p300 and H3K4me1 (Fig. S8). Together, these data suggest that PAXIP1 co-localizes with cohesin particularly at active promoters and enhancers.

## DISCUSSION

Here we present a high-confidence network of genetic dependencies of DDX11 and ESCO2 deficient cells. This reveals multiple genes previously linked to sister chromatid cohesion that confirm the presence of parallel cohesion establishment pathways ^10, 11, 23^. In addition, we validated the microtubule motor protein DYNC1Li2 as one of many mitotic regulators that become particularly essential in a cohesion-compromised background, thus representing potential therapeutic targets for cohesion defective cancers ^22^. Confidence of the identified hits is further underscored by multiple previously described interactions with genes involved in the response to DNA damage and DNA replication stress. While we chose to further investigate the PAXIP1-PAGR1 complex, it will be interesting to experimentally follow up some of the other newly identified genetic interactions.

We find that PAXIP1-PAGR1 directly promotes the chromatin association of cohesin. Our ChIP-seq analyses suggest that PAXIP1 particularly co-localizes with cohesin on sites that are not bound by CTCF. While CTCF-bound cohesin sites are mostly similar in different tissues, non-CTCF binding cohesin often localizes to cell-type specific transcription factors and active enhancers in specific genomic regions, frequently associated with cell identity genes ^66-70^. Since PAXIP1 binding is also enriched at promoters and enhancers and is necessary for long-range enhancer-promotor contacts ^45, 65, 71^, a function that is shared with cohesin ^36, 72, 73^, PAXIP1 may facilitate enhancer-promotor contacts by cohesin mediated loop formation. In line with this model, cohesin binding to glucocorticoid receptor (GR) binding sites depends on PAXIP1, resulting in a joint regulation of hormone-induced transcriptional activity via chromosome folding, described in the accompanying paper ^74^. Together, a picture emerges in which PAXIP1 facilitates chromatin binding of cohesin to regulate enhancer-promoter interactions, to control cell-type specific and context-responsive gene expression patterns.

Several mechanisms may explain how PAXIP1 promotes the chromatin association of cohesin. Cohesin and the cohesin loader co-precipitated with PAXIP1, similar as in other cellular models ^54, 62^. Notably, these interactions were difficult to detect, possibly indicating that they are weak, transient and/or context-dependent. Thus, PAXIP1-PAGR1 may act as a direct chromatin acceptor for the cohesin loading reaction, as has been suggested for several other chromatin binding proteins ^29, 30, 35, 75-77^, or stabilize cohesin on chromatin. This is further supported by FRAP analysis of SMC1-EGFP, which revealed reduced cohesin stability on chromatin in PAXIP1-KO cells ^74^. This may involve the FDF motif that we identified in PAGR1, which may antagonize WAPL similar as has been shown for CTCF ^28^ and MCM3 ^61^. The observation that PAXIP1 co-localizes with cohesin mainly at sites not occupied by CTCF could be consistent with the fact that cohesin can only bind one F/YxF motif at a time ^28^. Alternatively, and not mutually exclusive, PAXIP1 may modulate chromatin to create an environment that promotes cohesin loading, for example by creating a nucleosome free template ^29^. Thus, PAXIP1-PAGR1 may promote recruitment and/or stabilization of cohesin onto chromatin, resulting in local cohesin enrichment.

PAXIP1 and PAGR1 are components of KMT2C/D H3K4 methyltransferase complexes, but also function as an independent sub-complex at multiple genomic loci ^46^. Our findings suggest that human KMT2C and KMT2D are not involved in promoting cohesin occupancy on chromatin, although we cannot exclude that incomplete depletions preclude a detectable effect. Of note, PAXIP1 functions together with cohesin in glucocorticoid receptor activity apparently independent from KMT2D/C ^74^, indeed pointing at a function of PAXIP1-PAGR1 separate from KMT2C/D in cohesin regulation. Nevertheless, two PAXIP1 mutants previously described to abrogate KMT2C/D binding ^46, 55^ were unable to restore chromatin-bound cohesin, suggesting that BRCT5-6 has roles other than KMT2C/D binding.

In addition to regulating gene transcription, PAXIP1 has been implicated in immunoglobulin class switching and V(D)J recombination in B cells and in T cell receptor recombination ^45, 46, 65^, processes that also depend on chromatin looping and cohesin ^71, 78-81^. Moreover, like cohesin, PAXIP1 localizes to DNA damage sites ^55, 82-85^ and PAXIP1 has been shown to be important for downstream cohesin functions at DSBs ^48^. Considering the apparent overlap of processes that are controlled by PAXIP1 and cohesin, it will be interesting to determine to what extent these processes involve a direct effect of PAXIP1 on cohesin.

Although PAXIP1 is a hit in both screens, the effects of PAXIP1 loss are particularly pronounced in ESCO2 mutant cells, which may be related to the different functions of DDX11 and ESCO2 in cohesion establishment. Possibly, cohesin complexes that are loaded in a PAXIP1-dependent fashion in G1 are converted to cohesive cohesin in S-phase, which would place PAXIP1 in the cohesin conversion pathway like DDX11 ^12^. Alternatively, since PAXIP1 was also shown to localize to the replication fork ^86^, it may assist in cohesion establishment directly at the fork. However, while we initially identified PAXIP1 for its function in sister chromatid cohesion in ESCO2-mut cells, PAXIP1-KO cells do not harbor cohesion defects. This is reminiscent of ESCO1 depletion which does not cause pronounced cohesion defects, except in the context of ESCO2 loss ^13, 15, 38^. Similar to PAXIP1 loss, a reduction of functional MAU2 or NIPBL reduces chromatin-bound cohesin but does not cause pronounced sister chromatid cohesion defects. Since PAXIP1 promotes cohesin association to chromatin during G1, this together suggests that the main cohesin function of PAXIP1 is regulating the role of cohesin in intra-chromosomal DNA-DNA contacts.

## MATERIAL AND METHODS

### Cell culture

All cell lines were maintained in Dulbecco’s Modified Eagles Medium (DMEM, Gibco) with 9% FCS, 1mM Sodium Pyruvate (Gibco) and Penicillin Streptomycin (Gibco). RPE1-hTert-TetonCas9-PuroKO-TP53KO cells, referred to as RPE1-WT throughout this manuscript, were described previously ^37^. HCT116-KMT2D-KO cells were a kind gift from Yiping He ^57^.

### CRISPR-Cas9 gene editing

For CRISPR-Cas9 based gene editing approaches, we used an inducible Tet-On Cas9 expression system in combination with transfection of synthetic crRNA (IDT). In short, Cas9 expression was induced by 200 ng/mL doxycycline followed by transfection with 20 nM equimolar crRNA:tracrRNA duplexes with 1:1000 RNAiMax (Life Technologies). Genomic DNA was isolated with direct PCR lysis reagent (Viagen Biotech) with Proteinase K (O/N 55 °C). Proteinase K was inactivated (20 min 82 °C) and the crRNA target site was amplified with One Taq Hot Start DNA polymerase kit (NEB), followed by Sanger sequencing (primers in Table S3). Gene editing efficiencies were assessed using the Synthego ICE analysis tool ^87^.

### Genome-wide CRISPR-Cas9 screens

Genome-wide CRISPR-Cas9 screens were performed with the TKOv3 library ^88^ in triplicate at 400-fold library representation as previously described ^37^. In short, cells were transduced at MOI 0.2 with lentiviral pLCKO-TKOv3 and 8 µg/mL polybrene, and selected for viral integration with 5 µg/mL puromycin for three days. Cells were then harvested to take a t=0 sample and reseeded with 200 ng/mL doxycycline to induce Cas9 expression. After every 3 population doublings, cells were passaged for a total of 12 population doublings with doxycycline. Genomic DNA was isolated using the Blood and Cell Culture DNA Maxi Kit (Qiagen) and integrated gRNA sequences were amplified by PCR using HiFi HotStart ReadyMix (KAPA). Resulting PCR products were used as a template in a second PCR reaction in which Illumina adapters and barcodes were added. Data analysis is extensively described elsewhere ^43^. Briefly, sequencing reads were mapped to the TKOv3 library sequences with no mismatch tolerance. End-point reads were normalized to t=0 values by multiplying the average t=0 count per guide by the t=12 / t=0 fold-change (pseudocount +1), and the normalized counts were used as input for DrugZ analysis.

### Flow cytometry based cell cycle assay

Cells were incubated for 10 min with 10 µM 5’-ethynyl-2’-deoxyuridine (EdU), harvested, fixed in 4% paraformaldehyde for 15 min and subsequently overnight in 70% EtOH at -20 °C. Cells were then permeabilized in 0.5% Triton X-100, blocked with 5% FCS and incubated with Histone H3 pS10 Alexa Fluor 647 in 1% BSA, followed by incubation for 30 min with Click-it reaction mixture (50 mM Tris-HCl pH 7.6, 150 mM NaCl, 4 mM CuSO_4_, 1 µM Picolyl Azide 5/6-FAM, 2 mg/mL Sodium-L-Ascorbate) for EdU detection. Cells were washed and resuspended in 1% BSA with DAPI and detected by flow cytometry on a BD LSRFORTESSA X-20 (BD Biosciences). Data analysis was done using FlowJo V10.

### Clonogenic and viability assays

Two days after transfection of crRNA:tracrRNA duplexes, 1000 cells/well were seeded in 6-wells plates for clonogenic assays and 500 cells/well in 96-wells plates for CellTiter-Blue assays. For clonogenic assays, cells were fixed in 100% ice-cold MeOH 10 days after crRNA transfection, followed by staining in 0.5% crystal violet with 20% MeOH. CellTiter-Blue assays were performed 7 days after transfection. After incubation with CellTiter-Blue reagent (Promega) for 4 h at 37 °C, fluorescence was measured at 560_Ex_/590_Em_ with an Infinite F200 microplate reader (Tecan). For Incucyte experiments, cell growth was monitored at a 4 hour interval by an Incucyte Zoom instrument (Essen Bioscience) with a 10x objective.

### Protein extractions and co-immunoprecipitations

RPE1 cells were lysed in buffer containing 50 mM Tris-HCl pH 7.4, 150 mM NaCl, 1% Triton X-100 and protease inhibitors (1h on ice). Alternatively, where indicated, separate chromatin-bound and soluble protein fractions were prepared. First, cells were lysed in lysis buffer 1 (50 mM Tris-HCl pH 7,5; 150 mM NaCl; 10% glycerol; 0,2% NP-40) for 10 min on ice and centrifuged at 1300g for 10 min. Supernatant was used as soluble fraction. The pellet was then washed twice, followed by incubation in lysis buffer 2 (50 mM Tris-HCl pH 7,5; 150 mM NaCl; 10% glycerol; 1,0% NP-40 + 5 mM MgCl2 + 5 units/µL Benzonase) for 2h on ice, centrifuged at max speed for 5 min and the supernatant was used as chromatin-bound fraction. For co-immunoprecipitation in RPE1 cells, Flag-tagged PAGR1 was precipitated using Protein A/G Plus agarose beads (Santa Cruz; SC2003) with mouse-anti-Flag antibody (M2, #3165, Sigma), and Venus-tagged PAXIP1 was precipitated using GFP-trap beads (Chromotek; gta-20) following the manufacturer’s protocol.

For co-immunoprecipitations in HEK293T, cells were collected in buffer A (10 mM HEPES pH 7.9, 1.5 mM MgCl2, 10 mM KCl, 340 mM sucrose, 10% glycerol, 0.1% Triton X-100, protease inhibitor, phosphatase inhibitor), vortexed and incubated for 5 minutes on ice. Nuclei were collected by centrifugation (1300 g for 4 minutes at 4°C) and resuspended in buffer B (3 mM EDTA, 0.2 mM EGTA, 1 mM DTT, protease inhibitor, phosphatase inhibitor). Chromatin was collected by centrifugation (1700 g for 5 minutes at 4°C) and lysed with lysis buffer (10 mM HEPES pH7.9, 3 mM MgCl2, 5 mM KCl, 140 mM NaCl, 0.1 mM EDTA, 0.5% NP-40, 0.5 mM DTT, protease inhibitor and 62.5 Units of benzonase (Sigma #E1014)). After incubation for 45 minutes at 4°C, chromatin lysate was collected after centrifugation (13200 g for 20 minutes at 4°C). 2 mg of chromatin lysate was rotated overnight at 4°C with 5 μg antibody: rabbit IgG (Diagenode #C15410206), anti-NIPBL (NIPBL#1, Bethyl #A301-779A, targeting a central region; NIPBL#2, a homemade antibody targeting the C-terminus), anti-SMC3 (homemade) or anti-PAXIP1 (Sigma #ABE1877). Protein-antibody complexes were precipitated with 10µL of protein G Dynabeads (Fisher #10003D), washed four times with lysis buffer and proteins were eluted by boiling in sample buffer for 5 minutes.

For western blots, proteins were separated using a 8-16% Mini-PROTEAN Precast Protein gels (BioRad) and transferred to Immobilon-P PVDF membrane (Millipore). After blocking in 5% dry milk in TBST-T, membranes were incubated in primary and subsequently secondary peroxidase conjugated antibodies (antibodies used are listed in Table S3). Protein bands were visualized by incubation with ECL prime (Amersham).

### Immunofluorescence

Cells were grown on coverslips, pre-extracted with 0.5% Triton X-100 (2 min RT) where indicated and fixed in 4% paraformaldehyde. After permeabilization in 0.3% Triton X-100, cells were blocked in blocking buffer (3% BSA, 0.3% Triton X-100 in PBS), incubated with the indicated antibodies diluted 1:500 in blocking buffer, washed three times with PBS and incubated with appropriate anti-mouse/rabbit Cy3 conjugated antibody. After washing with PBS 3 times, coverslips were mounted with DAPI gold antifade (Invitrogen). Images were acquired using fluorescence microscopy (Leica). Analysis was performed using ImageJ. Background was subtracted using rolling ball background subtraction before quantification of nuclear intensity.

### Cohesion defect analysis

Cells were treated with 200 ng/mL democolcin for 20 min before harvesting and were subsequently swollen in 0.075 mM KCl. Next, cells were fixed in 3:1 methanol:acetic acid, dropped onto glass slides and stained in 5% Giemsa solution. For each condition 50 metaphases from coded slides were assessed for railroad chromosomes and premature chromatid separation. Metaphases were scored as normal with 0-4 railroad chromosomes, as partially separated with 5-10 railroad chromosomes and as single chromatids with at least two chromosomes completely separated.

### siRNA experiments

For siRNA experiments, cells were transfected with 20 nM siRNA using 1:1000 RNAiMAX and analyzed after 48h, unless otherwise stated.

### Lentiviral constructs and transduction

DDX11 and empty vector expression construct were previously described ^19^. To generate PAXIP1 and ESCO2 expression constructs, cDNA from RPE1 cells was cloned into pLenti CMVie-IRES-BlastR (Addgene plasmid #119863). PAXIP1 W75R, W175R and W676A mutants were constructed using overlap extension PCR. PAGR1 cDNA from HEK293T cells was N-terminally tagged with FLAG and also cloned into pLenti CMVie-IRES-BlastR. Lentiviral particles were produced in HEK293T cells and transduced into the indicated RPE1 cell lines, followed by selection with 10 µg/ml blasticidin or 5 µg/ml puromycin (Invitrogen).

### RNA-seq

Total RNA was isolated using the RNeasy mini kit (Qiagen). Up to 5 × 10^6^ cells per sample were lysed in RLT buffer. Samples were enriched for mRNA using the KAPA mRNA Hyperprep kit (Roche) and prepared for sequencing using the TruSeq RNA Library Prep Kit v2 (Illumina) according to the manufacturer’s instructions, and sequenced on an Illumina HiSeq4000. The Fastq files were clipped and cleaned by fastp ^89^. The clipped Fastqs were mapped to the human reference genome (hg19) by HISAT2 alignment tool ^90^. SAM to BAM transformation as well as sorting and indexing were carried out by SAMtools ^91^. Subsequently, the gene-level raw reads were counted by Subread’s featureCounts ^92^. Multi-mapping reads were also counted via assigning fractional counting to the genes. The raw count matrix was normalized and the differential expression analysis was performed by edgeR ^93^. The original library size was normalized to the effective library size by trimmed mean of M-value (TMM), followed by estimating dispersion by fitting the generalized linear model (GLM) with the design matrix. Subsequently, likelihood ratio test was performed to examine the differential expressions between WT and PAXIP1-KO samples. The differential expressions with false discovery rate (FDR) < 0.05 were defined as significant.

### ENCODE ChIP-seq data analysis

DeepTools (Version 3.5.1) ^94^ was used to generate heatmaps of publicly available ENCODE ChIP-seq data from HepG2 cells. Regions with peak summits (bed files) for RAD21 (ENCODE accession ENCFF052UCF), PAXIP1 (ENCFF825TUP), or POLR2A (ENCFF354VWZ) were sorted on signal (bigWig file) for RAD21 (ENCFF047SRI), PAXIP1 (ENCFF080TDD), or POLR2A (ENCFF425QWO), respectively, in a 2kb window using computeMatrix (--referencePoint --referencePoint=center -- beforeRegionStartLength=1000 -- beforeRegionStartLength=1000 --sortRegions=descend). Similarly, to look at active enhancers versus promoters, regions with p300 (EP300) peaks (ENCFF827LSX) were sorted on H3K4me1 signal (ENCFF470RYT) in a 4kb window. Sorted bed files for RAD21, PAXIP1, POLR2A, or p300 were used to generate matrices with bigWig files from ChIP-seq signals for CTCF (ENCFF938HDS), p300 (ENCFF962CGI), H3K4me3 (ENCFF359LQU) and/or H3K4me1 (ENCFF470RYT) using computeMatrix (-- sortRegions=keep). Heatmaps were generated using the plotHeatmap tool (--sortUsing=mean -- sortRegions=keep).

## AVAILABILITY

RNA-seq data are deposited at the National Center for Biotechnology Information (NCBI) Gene Expression Omnibus (GEO), accession number GSE211349. The datasets of ChIP-seq experiments analyzed in this study are from ENCODE, details are provided in the Methods section.

## ACKNOWLEDGEMENTS

We thank Isabel Mayayo Peralta and Wilbert Zwart for sharing unpublished work and for insightful discussions, the ENCODE Consortium and the Bernstein (Broad) and Myers (HAIB) lab for generating ENCODE datasets, and Yiping He for sharing HCT116 KMT2D-KO cells.

## Supplemental Figures 1-8

**Figure S1.**
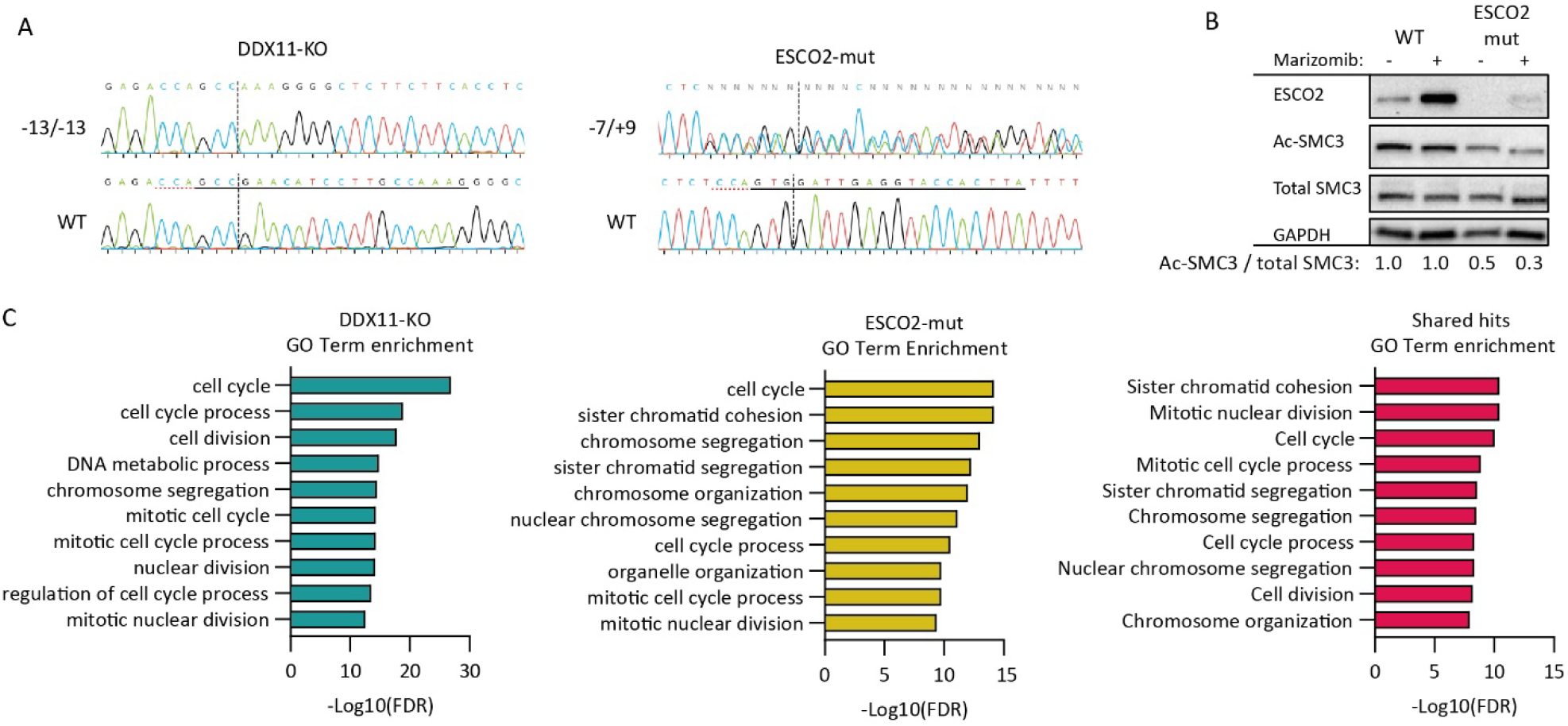
Genome-wide CRISPR screening in cohesion defective RPE1 cells. (**A**) Sanger sequencing of DDX11-KO and ESCO2-mut cells at the respective crRNA target sites. Resulting indels are indicated. (**B**) Western blot of WT and ESCO2-mut cells after treatment with 500 nM Marizomib for 5 h to inhibit protein degradation. Note the stabilization of residual ESCO2 protein after Marizomib treatment in ESCO2-mut cells. (**C**) Top 10 enriched gene ontology (GO) terms of the hits from DDX11-KO and ESCO2-mut screens, extracted from STRING-db.

**Figure S2.**
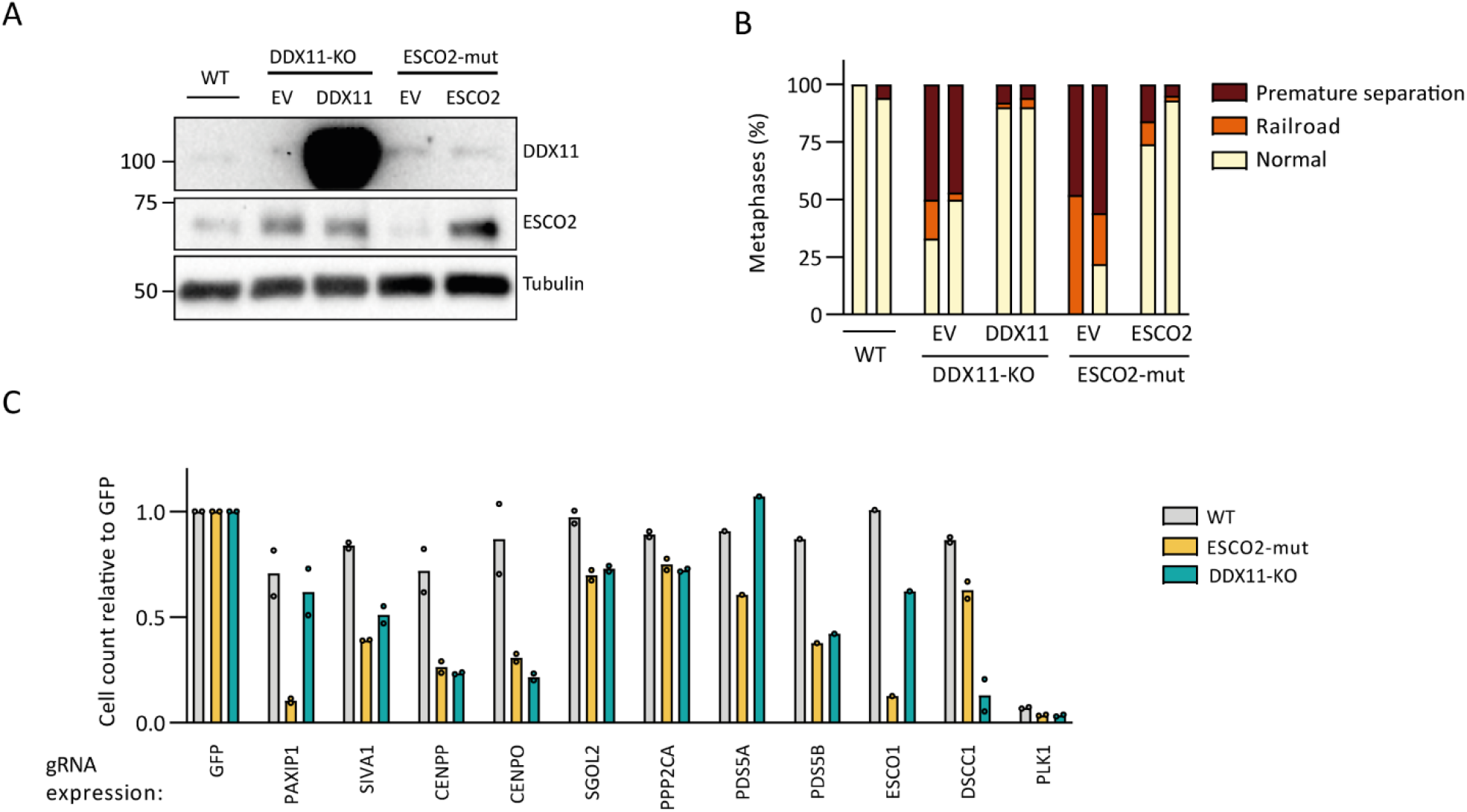
Extended validation of CRISPR screen results. (**A**) Western blot of ESCO2-mut and DDX11-KO cells with ectopic expression of empty vector (EV), DDX11 or ESCO2. (**B**) Cohesion defect analysis of ESCO2-mut and DDX11-KO cells with ectopic expression of EV, DDX11 or ESCO2. N=50 for each sample, two independent experiments are shown as separate bars. (**C**) Cells were stably transduced with indicated sgRNAs and cells were counted 10 days after Cas9 induction. Average viability is shown relative to sgGFP transduced cells; each dot represents an independent experiment.

**Figure S3.**
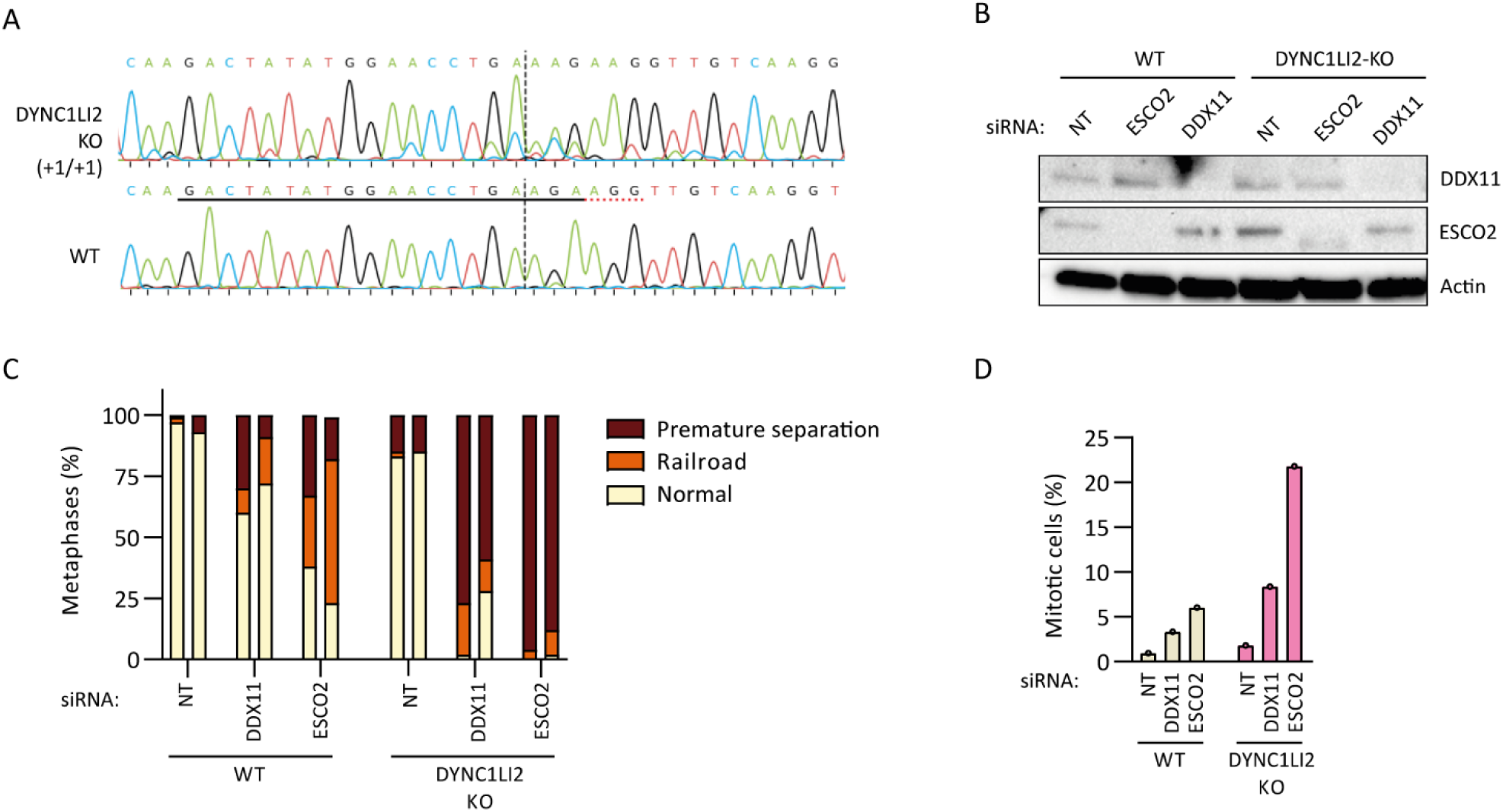
DYNC1Li2 is critical to mitigate further loss of cohesion in cells experiencing cohesion defects. (**A**) Sanger sequencing of DYNC1Li2-KO cells at the crDYNC1Li2 target site. Resulting indels are indicated between brackets. (**B**) Western blot control of indicated cell lines two days after transfection with indicated siRNAs. (**C**) Cohesion analysis of indicated cell lines two days after siRNA transfection. N=50 for each sample, two independent experiments are shown as separate bars. (**D**) Percentage of mitotic cells as assessed by flow cytometry of phospho-Histone H3 (Ser10) positive cells two days after siRNA transfection.

**Figure S4.**
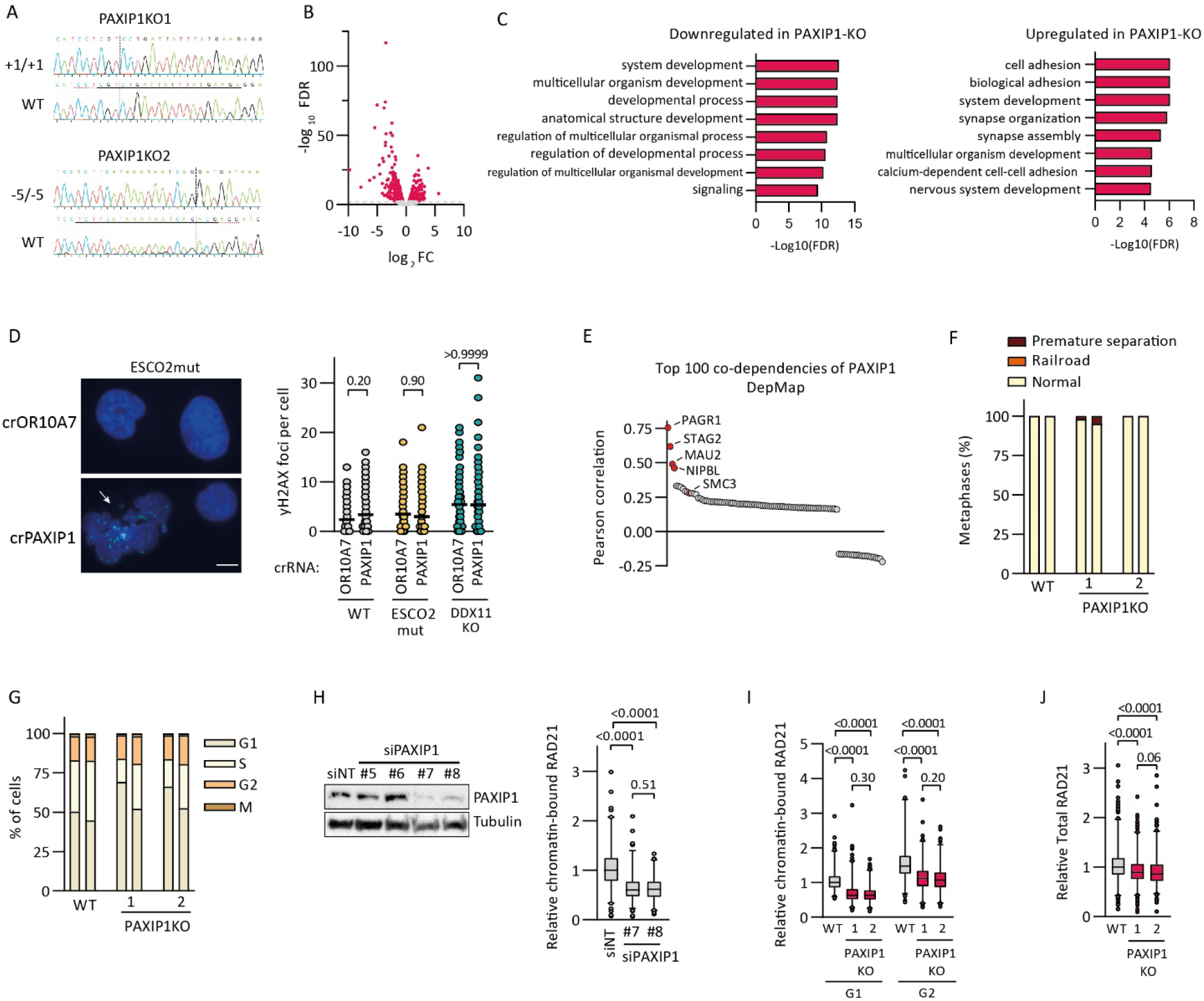
PAXIP1 promotes chromatin association of cohesin throughout the cell cycle independent of transcriptional changes. (**A**) Sanger sequencing of WT, PAXIP1-KO1 and PAXIP1-KO2 cell lines at the crPAXIP1 target site. Resulting indels are indicated. (**B**) Volcano plot of RNA-Seq of PAXIP1-KO versus WT. Negative log_2_ Fold Change (FC) indicates lower expression in PAXIP1-KO cells. 664 genes are significantly changed in expression (-log_10_ FDR > 2), depicted as red dots. (**C**) Top 8 Gene Ontology (GO) terms extracted from STRING-db of 357 genes downregulated (left) and 307 genes upregulated (right) in PAXIP1-KO versus WT. (**D**) γH2AX foci quantification two days after crPAXIP1 transfection in Cas9 expressing cells. At least 42 cells per sample were scored in two independent experiments. Only non-apoptotic, non-mitotic cells (assessed by circularity of intact nuclei) were scored. The arrow indicates a representative cell with disrupted nuclear integrity, likely resulting from mitotic catastrophe. Scale bar represents 10 µm. P-values were calculated by a one-way ANOVA. (**E**) Pearson correlation of the top 100 PAXIP1 co-dependencies based on the CRISPR Avana database from the DepMap portal (https://depmap.org). (**F**) Cohesion defect analysis of WT and PAXIP1-KO cells. N=50 for each sample, two independent experiments are shown as separate bars. (**G**) Cell cycle distribution of WT and PAXIP1-KO cells as assessed by DAPI, EdU incorporation (S-phase) and p-Histone H3-S10 (mitosis) using flow cytometry, in two independent experiments. (**H**) Cells were transfected with indicated siRNAs and analyzed by immunofluorescence with pre-extraction, to assess chromatin-bound cohesin levels (right). Data shown are from two independent experiments with >76 cells analyzed per sample. P-values were calculated by a one-way ANOVA. Box represents 25%-75% and median, whiskers represent 1-99 percentile. The western blot shows a PAXIP1 knockdown control (left). (**I**) Chromatin-bound cohesin levels of WT, PAXIP1-KO1 and PAXIP1-KO2 cells assessed by RAD21 immunofluorescence of pre-extracted cells synchronized in G1 or G2 by 24 h treatment with 10uM Palbociclib or 10uM RO-3306, respectively. RAD21 intensity quantified from three independent experiments with at least 137 cells per sample. P-values were calculated by a one-way ANOVA. Box represents 25%-75% and median, whiskers represent 1-99 percentile. (**J**) Total RAD21 levels of WT and PAXIP1-KO cells as assessed by RAD21 immunofluorescence without pre-extraction. At least 372 cells per sample were quantified in three independent experiments. P-values were calculated with a one-way ANOVA. Box represents 25%-75% and median, whiskers represent 1-99 percentile.

**Figure S5.**
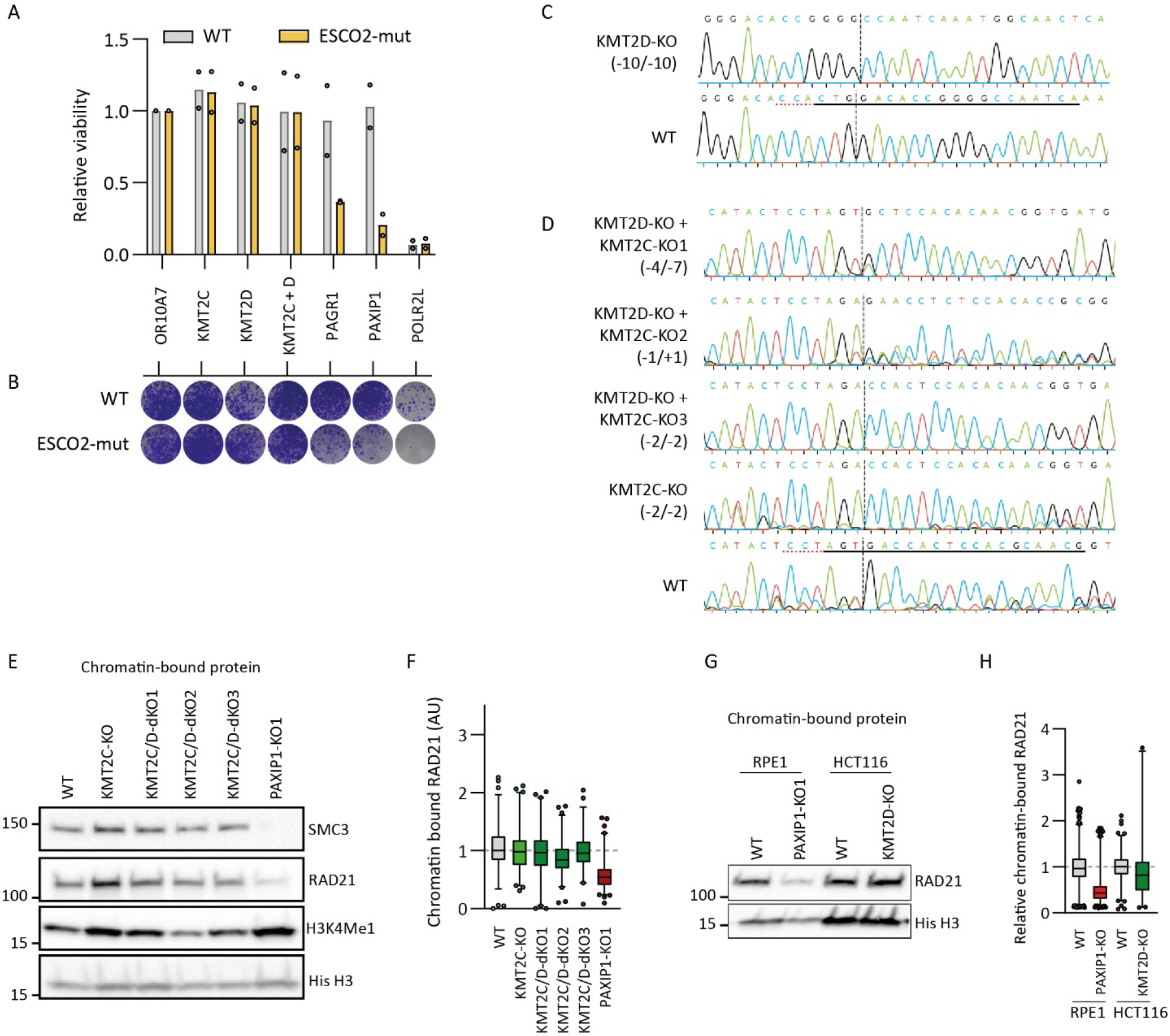
No evidence for involvement of histone methyltransferases KMT2C/D in promoting cohesin levels on DNA. (**A**) Viability of WT and ESCO2-mut cells assessed by a CellTiter-Blue assay 7 days after crRNA:tracrRNA transfection and accompanying Cas9 induction. Viability is normalized to cells transfected with crOR10A7, a nonessential gene. POLR2L is used as a common essential control. Dots represent the mean of three technical replicates; bars indicate the mean of two independent experiments. (**B**) Clonogenic survival assay 10 days after crRNA:tracrRNA transfection in indicated cells and accompanying Cas9 induction. (**C**) Sanger sequencing of KMT2D-KO clone at the crKMT2D target site. Resulting indels are indicated within brackets. (D) Sanger sequencing of KMT2C-KO and KMT2C/D-dKO clones at the crKMT2C target site. Resulting indels are indicated within brackets. (**E**) Western blot of chromatin-bound protein fractions. (**F**) Chromatin-bound cohesin levels assessed by RAD21 immunofluorescence of pre-extracted cells. RAD21 intensity from two independent experiments with at least 97 cells per sample are quantified. (**G**) Chromatin-bound cohesin levels of WT and PAXIP1-KO1 RPE1 cells and WT and KMT2D-KO HCT116 cells. Note that HCT116 cells already have homozygous loss of KMT2C. (**H**) Chromatin-bound cohesion levels assessed by RAD21 immunofluorescence of pre-extracted cells. RAD21 intensity from two independent experiments with at least 62 cells per sample are quantified.

**Figure S6.**
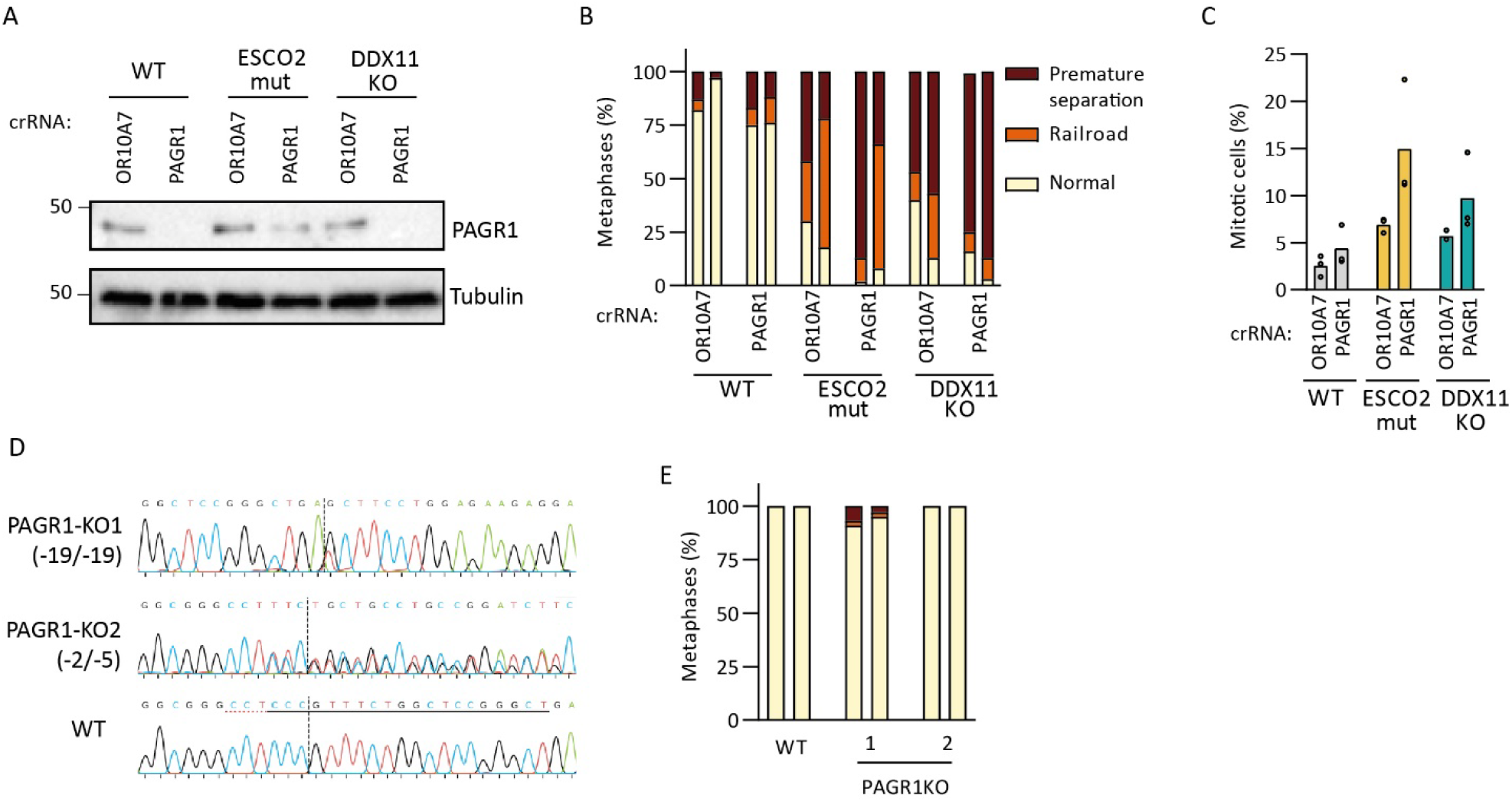
PAGR1 promotes functional cohesion in a sensitized background. (**A**) Western blot of whole cell extract of crOR10A7 or crPAGR1 transfected WT, ESCO2-mut and DDX11-KO cells. (**B**) Cohesion defect analysis two days after crOR10A7 or crPAGR1 transfection. N=50 for each sample, two independent experiments are shown as separate bars. (**C**) Percentage of mitotic (Histone H3 Phospho-Ser10 positive) cells two days after crPAXIP1 transfection from three independent experiments. (**D**) Sequences of indicated PAGR1-KO clones at crPAGR1 target site in exon 3 (at codon 196 of 255). Dotted line indicates Cas9 cut site. Resulting indels are indicated between brackets. (**E**) Cohesion analysis of WT and PAGR1-KO cells. N=50 for each sample, two independent experiments are shown as separate bars.

**Figure S7.**
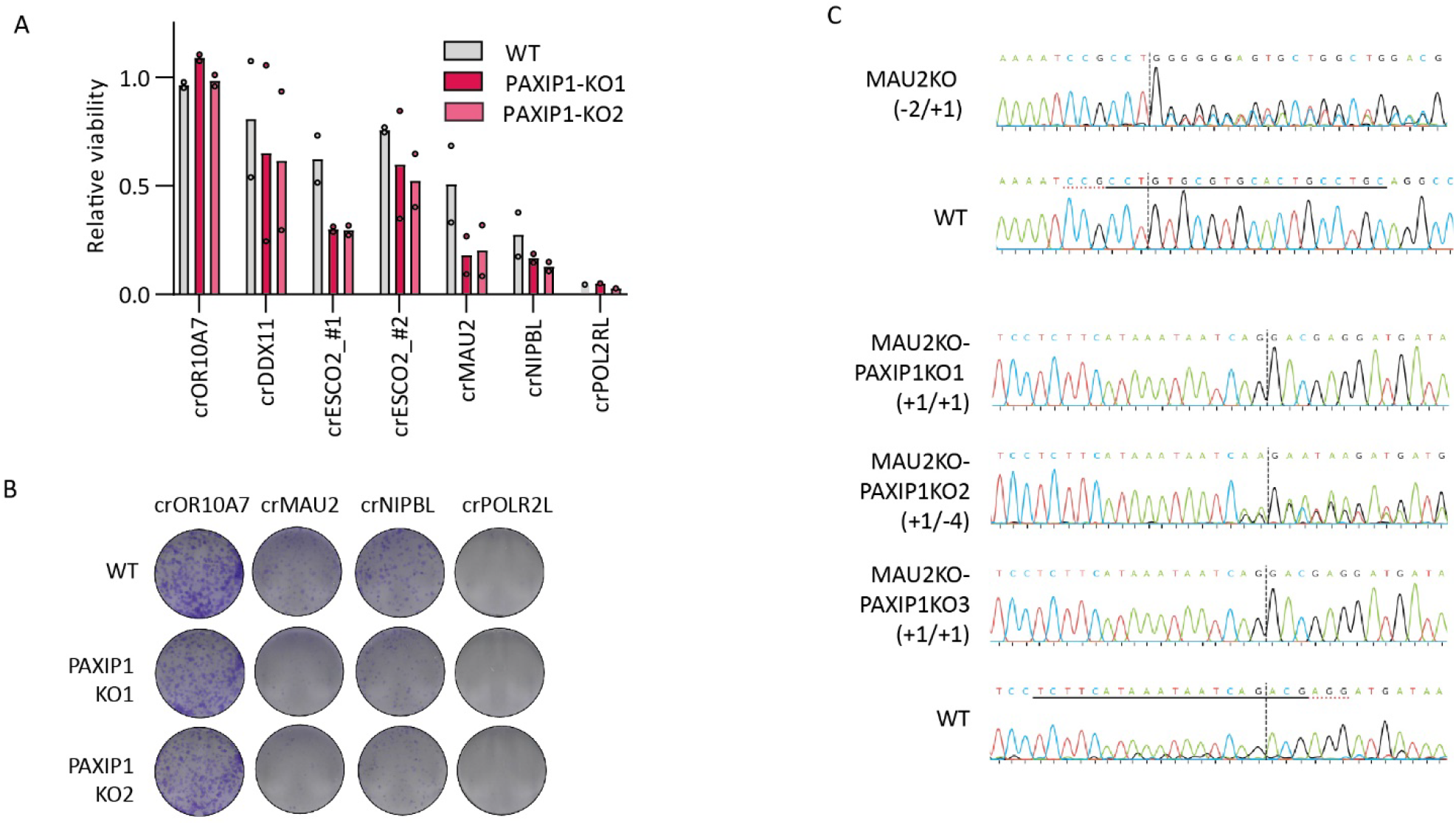
Increased sensitivity of PAXIP1-KO clones to depletion of NIPBL and MAU2. (**A**) CTB viability assay 7 days after transfection with indicated crRNAs in WT and PAXIP1-KO cells. Graphs represent mean values from two experiments using three technical replicates per experiment. (**B**) Clonogenic assay 10 days after transfection with indicated crRNAs in WT and PAXIP1-KO cells. (**C**) Sequences of indicated KO clones at crRNA target sites.

**Figure S8.**
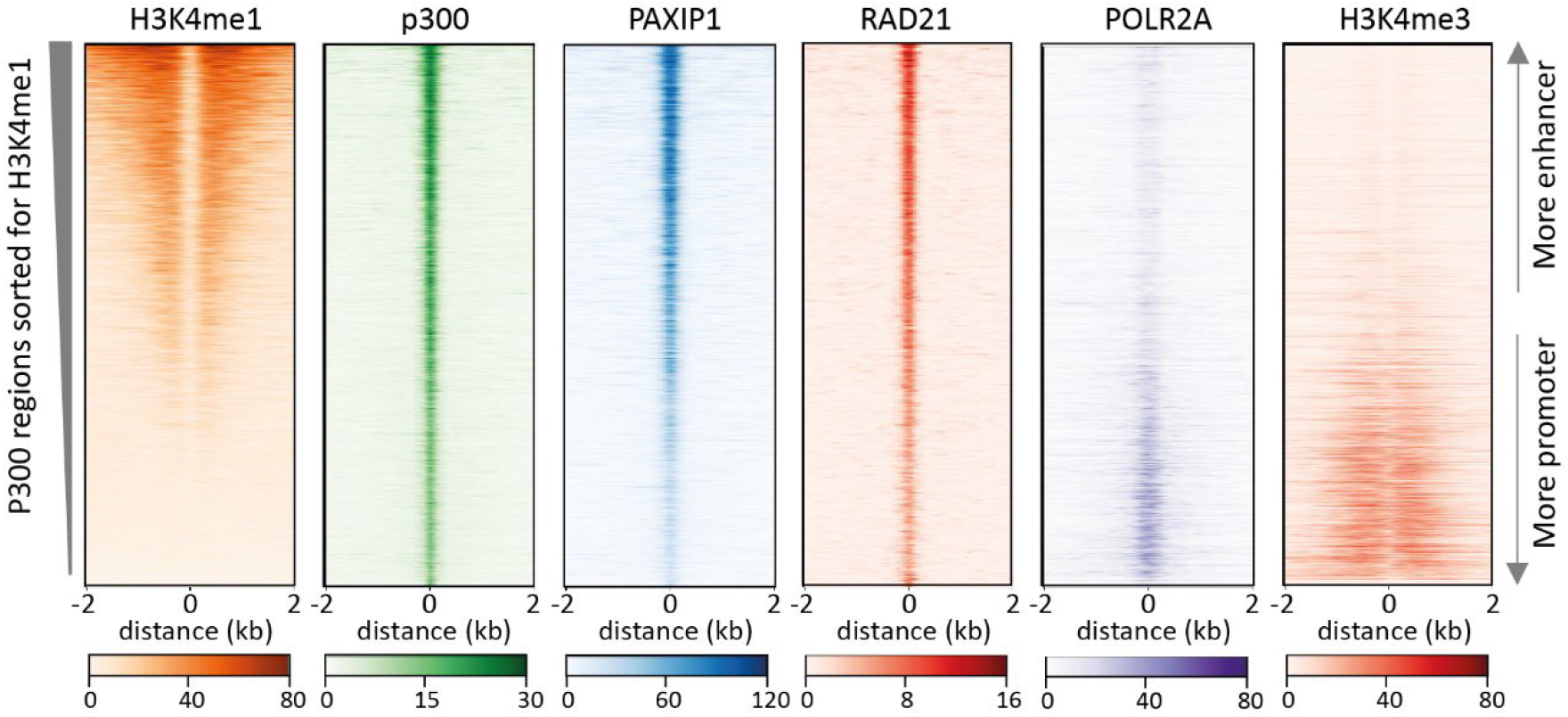
PAXIP1 co-localizes with cohesin on active promoters and enhancers. Heatmaps of ENCODE ChIP-seq data for HepG2 cells. Regions with peaks for p300 were sorted for H3K4me1 signal.

